# Human TDP-43 overexpression in zebrafish motor neurons triggers MND-like phenotypes through gain-of-function mechanism

**DOI:** 10.1101/2025.07.06.663393

**Authors:** Alison L Hogan, Madison Kane, Patrick Chiu, Grant Richter, Cindy Maurel, Sharlynn Wu, Natalie M Scherer, Emily K Don, Albert Lee, Ian Blair, Roger Chung, Marco Morsch

## Abstract

Dysregulation of the TAR DNA-binding protein 43 (TDP-43), including intraneuronal cytoplasmic mislocalisation and aggregation is a feature of multiple neurodegenerative diseases including amyotrophic lateral sclerosis (ALS), frontotemporal lobar dementia (FTLD), limbic-predominant age-related TDP-43 encephalopathy (LATE) and alzheimers disease (AD). Unravelling the causes and functional consequences of TDP-43 dysregulation is paramount to understanding disease mechanisms as well as identifying effective therapeutic targets. Here we present a comprehensive *in vivo* characterisation of three stable transgenic zebrafish models that express human TDP-43 variants in motor neurons. We demonstrate that overexpression of predominantly nuclear wildtype TDP-43, cytoplasm-targeted TDP-43, and an ALS-linked variant (G294V) each induce toxic gain-of-function effects, leading to impaired motor function, motor neuron loss, and muscle atrophy. Importantly, these models reveal distinct phenotypes, with the ALS-linked mutant exhibiting axonal transport deficits and neuromuscular junction disruption, while cytoplasmic mislocalised TDP-43 heightened susceptibility to oxidative stress. Two FDA-approved drugs used to treat ALS, edaravone and riluzole, were examined in these models and revealed that edaravone, but not riluzole, was effective in rescuing motor deficits associated with cytoplasmic TDP-43 expression and, to a lesser extent, mutant TDP-43^G294V^. Collectively, these findings reveal distinct pathological consequences of TDP-43 dysregulation, providing neuron-centric mechanistic insights, and establish the humanised TDP-43 zebrafish as an efficient system for preclinical therapeutic testing.

**Graphical abstract:** 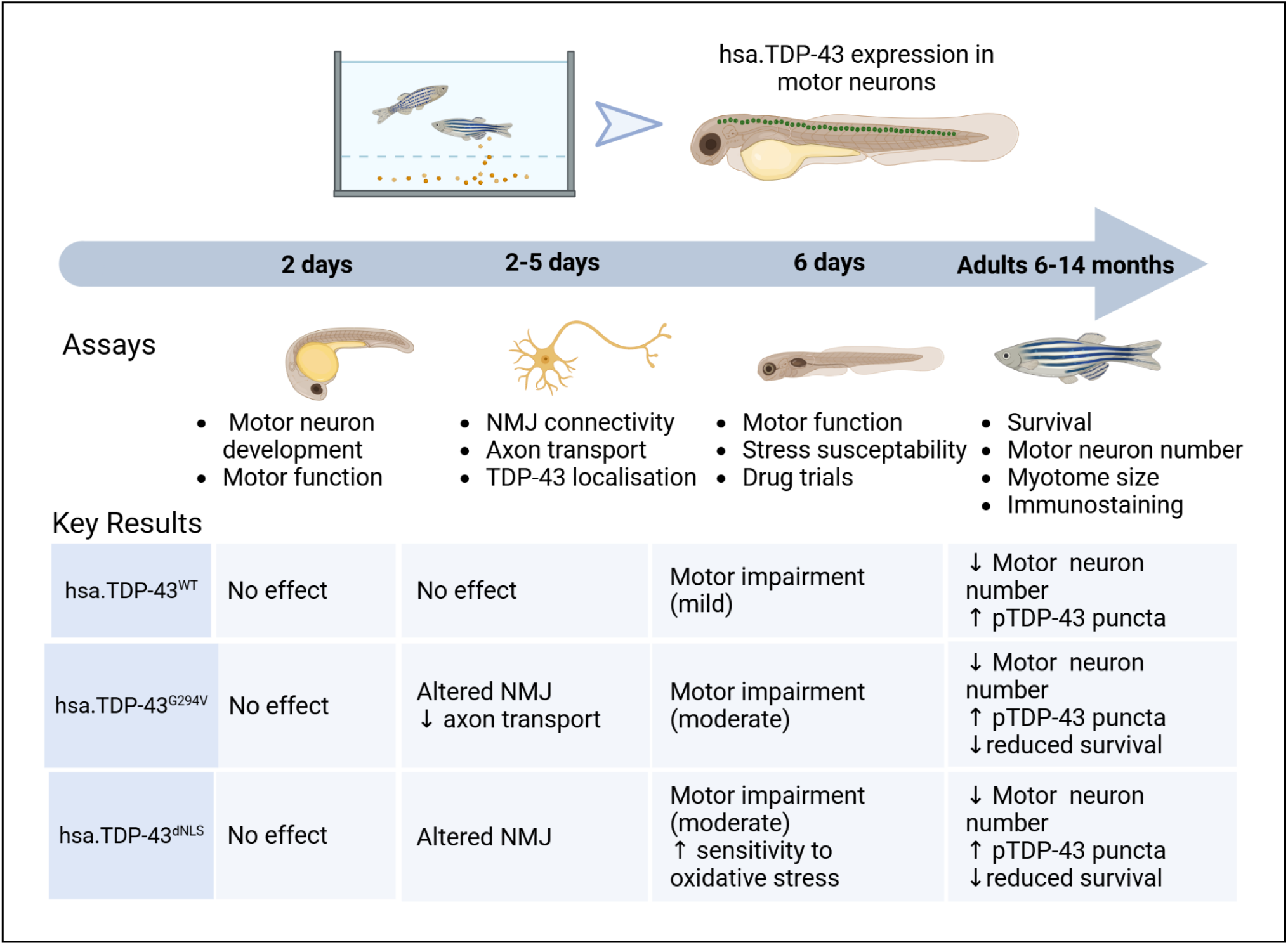

## Introduction

TAR DNA-binding protein 43 (TDP-43) is a DNA and RNA-binding protein that plays a central role in the pathogenesis of several neurodegenerative diseases [1]. Cytoplasmic mislocalisation of this predominantly nuclear protein, along with its hyperphosphorylation, ubiquitination and accumulation in insoluble aggregates, are neuropathological hallmarks of amyotrophic lateral sclerosis (ALS) [2], [3], frontotemporal lobar dementia (FTLD) [3], and limbic-predominant age-related TDP-43 encephalopathy (LATE) [4]. Cytoplasmic mislocalisation is also observed in up to 57% of Alzheimer’s disease (AD) patients [5]. In addition, mutations that cause ALS [6], [7] and FTLD [8] have been identified, across all functional domains, in the gene that encodes TDP-43, *TARDBP*. While mutations in *TARDBP* are a rare cause of ALD and FTLD (<5% of patients [9]), the commonality of TDP-43 pathology across ALS (>90% of patients [10]), FTLD (≤ 50% of patients [3]), LATE (100% of patients [4]) and AD (∼57% of patients [5]) underscores the central role of TDP-43 in the pathogenesis of these diseases. Notably, while TDP-43 pathology is a common pathological hallmark for these diseases, the underlying molecular mechanisms are likely to be diverse.

No model system recapitulates ALS, FTLD, LATE, or AD in all facets [11]. However, they do provide valuable tools to study specific aspects of the mechanistic cascade of TDP-43 pathology, with numerous *in vitro* and *in vivo* models developed for this purpose (reviewed in [12]–[15]). Reported vertebrate models include those that display overexpression of wildtype TDP-43 [16]–[19], TDP-43 depletion [20]–[22], TDP-43 aggregation [23], TDP-43 with ALS-linked mutations (overexpressed and at endogenous levels) [24]–[31], models with cytoplasmic accumulation and concomitant nuclear depletion of TDP-43 though CRISPR-directed genome editing [32], or overexpression of a TDP-43 transgene with a disrupted nuclear localisation sequence (dNLS) [33], [34]. Functional studies of these models, combined with *in vitro* studies, have identified a wide array of mechanisms associated with TDP-43 related neurodegeneration. These include oxidative stress and hyper-responsiveness to cellular stress, mitochondrial dysfunction, altered axon transport, synaptic dysfunction and hyperexcitability (reviewed in [35]). Recent seminal studies have shown that the loss of nuclear TDP-43 function, acting as a repressor of cryptic exon retention, results in cryptic RNA splicing and subsequent downregulation of key proteins including STMN2 [36] and UNC13A [37].

Zebrafish have proven to be useful additions to the TDP-43 toolkit for study of these neurodegenerative diseases. The optical transparency of this model facilitates unparalleled real-time imaging of protein dynamics [38], [39], cell-cell interactions [40] and targeted manipulations, such as optogenetic induction of oxidative stress [41] or aggregation [23], [42] in a single neuron, studies that are significantly more challenging in other vertebrate models. Additionally, zebrafish models enable time-efficient testing of potential therapeutics *in vivo*.

We recently demonstrated the utility of zebrafish to study protein behaviour at the molecular level by overexpressing fluorescently tagged human TDP-43 (hsa.TDP-43) in individual zebrafish motor neurons. This demonstrated for the first time that hsa.TDP-43 undergoes liquid-liquid phase separation (LLPS) to form bimolecular condensates in spinal motor neurons *in vivo* and that RNA-binding deficiencies and post-translational modifications can alter this behaviour [38]. This has provided valuable insight into the dynamic nature of phase transitions of TDP-43 and the potential for functional, reversible bimolecular condensates to progress to disease-associated aggregates.

Here we present a comprehensive characterisation of three stable transgenic zebrafish lines that constitutively overexpress hsa.TDP-43 within motor neurons of the spinal cord. We generated zebrafish that express either wildtype hsa.TDP-43 (hsa.TDP-43^WT^), an ALS-causative mutation located within the C-terminal domain (hsa.TDP-43^G294V^), or hsa.TDP-43 with a disrupted nuclear localisation signal (TDP-43^dNLS^), to investigate the *in vivo* functional consequences. While embryonic development appeared largely unaltered in these models, a robust motor phenotype was observed by 6 days of age in hsa.TDP-43^G294V^ and hsa.TDP-43^dNLS^ zebrafish. The effects of hsa.TDP-43 on motor neuron health were also evidenced by progressive loss of motor neurons and reduced muscle mass with old age. Interestingly, differences in neuromuscular junction integrity, axonal transport, susceptibility to oxidative stress and response to drug treatments (riluzole and edaravone) were observed between the various hsa.TDP-43 expression systems, indicating distinct vulnerabilities and functional consequences.

Overall, the three zebrafish models provide unique insights into the mechanisms of TDP-43 pathology and demonstrate the suitability of these platforms as ‘testbeds’ for assessing novel therapeutic interventions. The observed phenotypes can be assessed as early as 6 days or long-term, providing a critical platform for accelerating clinical outcomes in the future.

## Materials and Methods

### Zebrafish husbandry

Zebrafish (*Danio rerio*) used in this study were maintained under standard conditions [43] including water temperature of 28°C, pH 7.4, 14 hour light-10 hour dark cycle and twice daily artemia and pellet feeds). Larvae were maintained in E3 embryo medium (5 mM NaCl, 0.17 mM KCl, 0.33 mM CaCl_2_, and 0.33 mM MgSO_4_ buffered to pH 7.3). Only morphologically normal embryos were used for analysis.

### Transgenic zebrafish used in the study

Zebrafish used in this study were AB/Tuebingen wild type lines that expressed EGFP-tagged wildtype human TDP-43 (hsa.TDP-43^WT^), cytoplasmic human TDP-43 with three amino acid substitutions (K82A, R83A, K84A) disrupting the Nuclear Localisation Signal (hsa.TDP-43^NLS^) or human TDP-43 with an ALS-causative mutation (hsa.TDP-43^G294V^) within motor neurons using the *-3mnx1* promoter. Generation of the constructs used to develop these lines has been previously described [38], [44]–[46].

For analysis of motor neuron axons, the hsa.TDP-43 transgenic lines were crossed with established transgenic lines that express blue fluorescent protein within motor neurons (Tg(*-3mnx1*:mtagBFP) [44].

### Western blot validation of hsa.TDP-43 expression

Zebrafish at 3 days post fertilisation (dpf) were anesthetized in 50 mg/L tricaine (MS222) on ice (n = 25-30). Embryos were deyolked in Ringer buffer (55 mm NaCl, 1.8 mm KCl, 1.25 mm NaHCO_3_) followed by PBS wash, then homogenised via pestle in RIPA buffer (20 mm Tris-HCl, pH 7.4, 1% Triton X-100, 150 mm NaCl, 1 mM EDTA and complete protease inhibitor cocktail, Roche Applied Sciences) followed by 20 minute agitation at 4°C and probe sonication. The homogenised sample was centrifuged for 20 minutes at 13,000 rpm at 4°C and the supernatant collected for analysis [38], [47].

Protein concentration was determined using the Pierce BCA Protein Assay Kit and western blotting performed as previously described [38]. Briefly, 30 µg protein lysate was loaded into a 4-15% SDS-PAGE gel (Biorad), electrophoresed for 40 minutes at 180V and transferred via semi-dry transfer onto a nitrocellulose membrane using a Bio-Rad Turbo Transfer apparatus (2.5A, 7 mins). Membranes were blocked in blocking buffer (PBS, 0.5% Tween 20, 3% BSA) for 1 hour at room temperature, followed by overnight incubation in primary antibody solution at 4°C. Primary antibodies used: TDP-43 rabbit polyclonal 1:10000 (Proteintech, 10782-2-AP), GAPDH rabbit polyclonal 1:15000 (Proteintech, 60004-1-Ig). Membranes were washed 3 times in PBS and 0.5% Tween 20 and then incubated for 45 minutes at room temperature with IRDye 800CW donkey anti-rabbit IgG and 680LT donkey anti-mouse IgG, 1:15000 (LCR-926-32212 and LCR-926-68023 respectively, LI-COR Biosciences). Membranes were visualised using the Odyssey CLx imaging system and protein bands quantified with the Image Studio Lite software (LI-COR).

### Analysis of motor neuron axon morphology

Zebrafish embryos at 48 hours post fertilisation (hpf) were manually dechorionated, anaesthetised with tricaine (MS222) and mounted in 3% methylcellulose. Imaging of BFP-tagged motor neurons was performed on a M165FC fluorescent stereomicroscope (Leica). The first 8 ventral axonal projections of the primary motor neurons immediately caudal to the yolk ball/tube boundary were assessed by 2 blinded researchers for the presence of abnormal branching. Branching at or above the ventral edge of the notochord or truncation of the axons was scored as aberrant [48], [49]. Average axonal length was also measured for the first five ventral axonal projections of this region using ImageJ Measure tool.

### Assessment of axonal transport

For analysis of axonal transport we generated a Tol2:*elavl*:mCherry_kif5a construct using the Multisite Gateway^®^ Three Fragment Vector Construction. The pTol2pA2 [50] destination vector was recombined with a p5E-*elavl3* promotor [44], pME-mCherry fluorescent tag (no stop codon) [44] and a p3E-DN-Kif5a vector, which was a gift from Caren Norden (Addgene plasmid # 105973; http://n2t.net/addgene:105973) [51]. Following injection of the Tol2:*elavl3:*mCherry_kif5a construct into eggs of hsa.TDP-43 zebrafish at 20 ng/µl in combination with transposase mRNA (30 ng/µL), embryos were screened for mosaic expression of mCherry-Kif5a and motor neuron expression of EGFP-hsa.TDP-43 at 48 hpf on a M165FC fluorescent stereomicroscope (Leica). At 3 dpf, double transgenic larvae were mounted in 1% low melting agarose and immersed in E3 embryo media with tricaine. mCherry-KIF5a positive axons were imaged on a Leica Stellaris 5 confocal microscope with a 25X water immersion lens and 8X magnification with bidirectional scanning speed of 700 and a small region of interest incorporating the axon only to minimise acquisition time, which ranged from 150 – 400 ms per image. Timelapse images were taken over a 5 minute period.

Analysis was performed using ImageJ. Raw images were processed by converting the colour scheme to “fire” and application of gaussian blur with a 0.2 µm sigma radius. Only puncta size 0.8 – 2 µm^2^ in size were analysed to enable confident tracking of a single particle and to minimise variability associated with large puncta size [52]. Movement of individual puncta (anterograde or retrograde) was tracked using the MtrackJ plugin. Puncta were tracked until they moved out of the plane of focus, out of frame, or merged with a second puncta and could no longer be tracked with confidence. A minimum of 50 particles from 5 zebrafish per group were analysed.

### Assessment of neuromuscular junctions

Immunohistochemistry was performed on 2- and 5-days post fertilisation (dpf) wild-type (WT), dNLS, and G294V TDP zebrafish to visualise presynaptic and postsynaptic structures at the neuromuscular junction (NMJ). Zebrafish larvae were fixed in 4% paraformaldehyde at 4 °C overnight. After rinsing with PBS-Tween, samples were blocked with 10% normal donkey serum in PBS-Tween 20. Larvae were incubated with alpha-bungarotoxin conjugate with Alexa Fluor™ 647 (10mg/mL; Invitrogen, B35450) in 0.1% PBS-Tween20 for 30 min. Larvae were then incubated overnight at 4 °C with the primary antibody SV2 (1:200, DSHB, University of Iowa, SV2-c). This was followed by a 2 hour incubation at room temperature with secondary antibody Alexa Fluor 488 (1:1000, Invitrogen). After washing, the larvae were mounted in 70% glycerol. Z-stack images were captured using a Leica Stellaris 5 confocal microscope with a 25X water immersion lens (Carl Zeiss, Germany) and processed with ZEN software (Carl Zeiss). Colocalisation of presynaptic and postsynaptic structures was quantified per somite using ImageJ software (NIH) from stacked Z-series images according to established protocols [53].

### Motor function assays and drug treatments

For all assessments of motor function, embryos were raised in equal numbers. For the 48 hpf photomotor response assay, embryos were raised in a dark incubator until 36 hours, then dechorinated and placed into a 96-well plate and returned to dark conditions. At 48 hpf, embryos were moved into a ZebraBox (Viewpoint), given 10 minutes to acclimatise (in the dark), then exposed to a 1 second, 300 watt light stimulus [54]. Distance swum in response to this stimulus was tracked using Zebralab software (View-point).

For the 6 dpf motor function assay, embryos were raised in equal numbers (n = 60 per dish) in a light-dark incubator. At 6 dpf they were moved to a 48-well plate, given an hour to recover, then moved into the ZebraBox (Viewpoint). After 10 minutes acclimatisation time, distance swum in alternating 5 minute light-dark cycles (20 minutes total) was recorded with Zebralab software (Viewpoint) [55].

For drug treatment assays, embryos were manually dechorionated at 2 dpf to ensure full exposure to drug treated E3 media. Drugs were replenished daily until tracking at 6 dpf. For assessment of susceptibility to oxidative stress, fresh 30% H_2_O_2_ was added to the E3 media from 3 dpf to a final concentration of 0.5 mM as previously optimised [56]. For Riluzole treatment, Riluzole (Cayman Chemical, cat#35833) suspended in DMSO was added to E3 media of dechorionated embryos at 3 dpf at the previously reported dose of 7.5 mM [57]. DMSO alone was added to the control / untreated group. The optimal dose of Edaravone in zebrafish has not been established in the literature, so a dose-response trial was first performed. Given their heightened susceptibility to oxidative stress, the hsa.TDP-43^dNLS^ zebrafish were used for this optimisation. Edaravone (Cayman Chemical Company, MCI-186) was dissolved in DMSO and added to the E3 media at concentrations of 100 μM, 40 μM, 20 μM, 10 μM and 5 μM. Doses >40 μM was found to be toxic (**Supplementary Fig 5**). Effects on motor function were therefore assessed only for 20 μM, 10 μM and 5 μM. A significant rescue effect was observed at 5 μM. 5 μm was therefore used for analysis.

### Adult zebrafish sectioning

Following euthanasia by submersion in ice cold water for 30 minutes, adult zebrafish at 6 months post fertilisation (mpf) and 14 mpf were segmented with a scalpel blade. The region directly ventral to the dorsal fin was collected and fixed in 4% paraformaldehyde in PBS overnight at 4°C. Following PBS washes, samples were dehydrated in 15%, then 30% sucrose in PBS overnight at 4°C. Dehydrated sections were then mounted in Optimal Cutting Temperature Compound (OCT, Fisher HealthCare) and frozen at −80 °C in preparation for cutting with a Leica CM1050 cryostat. For motor neuron counts, 30 µm thick sections were mounted on Superfrost plus slides (Thermofisher). For immunostaining, free-floating 50 µm sections were cut and stored in PBS with 0.03% sodium azide until staining.

### Nissl staining for motor neuron counts in adult zebrafish

Mounted sections were dried at room temperature (30 minutes), rehydrated in PBS (30 minutes), permeabilised in PBS plus 0.1% triton X-100 (10 minutes) then washed in PBS. Sections were then stained with NeuroTrace 640/660 (Thermofisher) at 1:150 dilution for 30 minutes. Following 3x PBS washes totalling 2 hours), sections were mounted with DAKO mounting medium and imaged. A minimum of 20 sections per zebrafish were imaged using a Leica Stellaris 5 confocal microscope. Z-stack images were taken, then combined into a maximum intensity projection image for analysis. ImageJ was used to identify Nissl-stained cells >75 μm^2^ in the ventral horn of the spinal cord as previously described [58].

### Analysis of myotome size

Mounted 30 µm thick sections of zebrafish at 14 mpf were also used for quantification of myotome size. To ensure the same muscle segments were examined for each fish, a region of interest was delineated by the dorsal and ventral borders of the spinal cord within 200 µm of the spinal cord. All myotomes within this region were used for analysis. Individual myotome size was determined using ImageJ Measure tool and mean myotome size determined for each zebrafish.

### Ageing assay

Zebrafish for this assay were raised under standard conditions. Clutches from all three lines from the F4 generation were collected on the same day and the same number of larvae housed in petri dishes (n = 30) until transfer to the system for feeding at 6 dpf. At one month of age, numbers were corrected to maintain equal housing density between groups and survival recording commenced. Zebrafish underwent daily health checks and any fish with anatomical abnormalities detected were euthanised by immersion in ice cold water a per following ethical guidelines. Survival was recorded monthly until 14 months.

### Immunofluorescent staining of adult zebrafish

Each strain group comprised three adult zebrafish aged to 14 months post-fertilisation. Zebrafish were euthanised, and tissues were dissected and sectioned to 55 µm using a cryostat immediately prior to staining.

Sections were permeabilised and blocked in 10% NGS in PBS for 2 hours at room temperature before incubation with primary antibodies targeting native TDP-43 (rabbit; 1:1000; Cat. No. 10782-2-AP) and phosphorylated TDP-43 (mouse; 1:500; Cat. No. TIP-PTD-M01) overnight at 4°C. After washing, sections were incubated with species-appropriate secondary antibodies (AlexaFluor 488 for pTDP-43 and AlexaFluor 555 for nTDP-43) for 2 hours at room temperature. Following secondary antibody incubation, sections were washed and counterstained with NeuroTrace 640/660 (Thermofisher) at 1:150 dilution for 30 minutes, then washed for an additional 2 hours. Slides were mounted using DAKO mounting medium containing DAPI and stored at 4°C until imaging.

### Statistical analysis

All analysis was replicated in zebrafish from a minimum of three clutches and only morphologically normal zebrafish were included in the study.

Statistical analysis was performed and graphs generated using GraphPad Prism 9 (9.2.0). Shapiro-Wilk normality test or the D’Agostino-Pearson normality test was used to assess normality distribution of data for each assay. Motor function assays were further examined for statistical outliers that may indicate inaccuracy for tracking using GraphPad Prism 9 ROUT method with Q value (false discovery rate) of 0.5%. Values determined to be outliers were excluded from analysis.

For normally distributed data, One-way ANOVAs and Holm-Sidak’s multiple comparison test were used to analyse datasets with >2 groups and Gaussian distribution and two-tailed t-tests for pairwise comparisons. For data with non-Gaussian distribution, Kruskal-Wallis test with multiple comparisons was used to examine significance. For all statistical tests significance was taken as *p < 0.05, **p<0.01, ***p < 0.001. Unless otherwise indicated, data values are presented as the mean ± standard error of the mean (SEM).

## Results

### Generation of stable transgenic zebrafish that overexpress human TDP-43 in motor neurons

To assess the consequences of stable motor-neuron expression of hsa.TDP-43 in the spinal cord, we generated transgenic zebrafish lines that express either wild-type human TDP-43 (hsa.TDP-43^WT^), an ALS-associated mutant (hsa.TDP-43^G294V^) or the mislocalised (cytoplasmic) variant hsa.TDP-43^dNLS^. All constructs were designed with a N-terminal green fluorescent protein (EGFP) and expression was driven by the *-3mnx1* motor neuron promoter (**Fig 1A**). Expression of the hsa.TDP-43 protein was confirmed in each transgenic line using fluorescent confocal microscopy from 2 dpf onwards and western blotting analysis at 3 dpf. Expression of the TDP-43 transgene throughout the spinal cord was observed for all transgenic lines. As previously reported [38], hsa.TDP-43^WT^ and hsa.TDP-43^G294V^ were predominantly expressed in the nucleus of motor neurons, while hsa.TDP-43^dNLS^ was primarily confined to the cytoplasm (**Fig 1B**). Western blot analysis at 3 dpf (n= 25–30 embryos per replicate, 3 replicates) demonstrated no significant difference in hsa.TDP-43 expression levels between transgenic lines when normalised to a GAPDH loading control (**Fig 1C, 1D,** full blot shown in **Supplementary Fig 1**).

**Figure 1:**
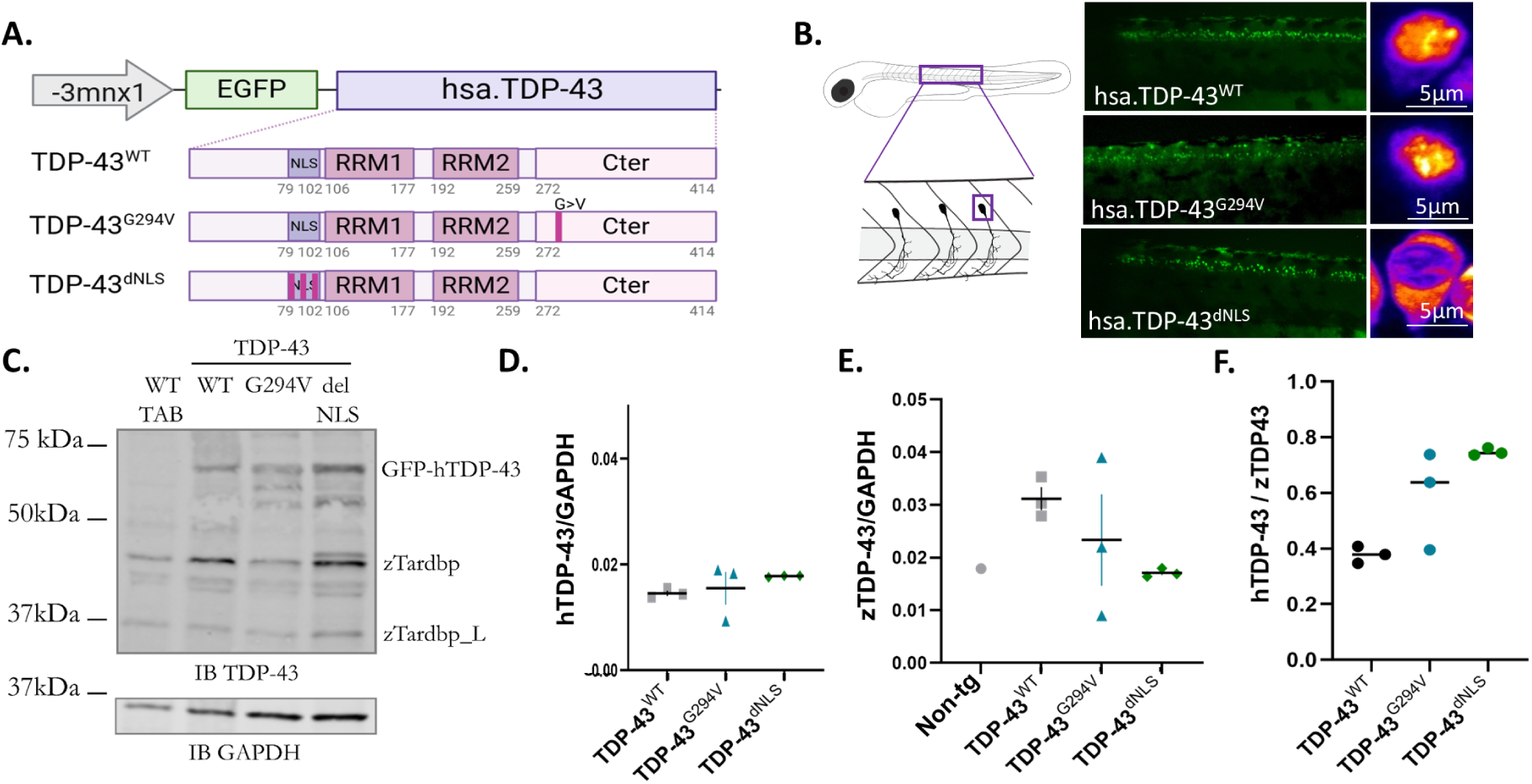
Validation of human TDP-43 (hsa.TDP-43) expression in zebrafish motor neurons. **A.** Schematic of constructs used to generate hsa. TDP-43 transgenic zebrafish. EGFP-tagged hsa.TDP-43 (wild-type, G294V or dNLS) was expressed under the −*3mnx7* neuronal promoter. **B.** Schematic and representative images showing the expression of hsa. TDP-43 along the spinal cord and within a single motor neuron soma for each transgenic line. **C.** Western blot analysis confirming expression of hGFP-TDP-43 protein. **D.** No significant difference in expression was observed between hsa. TDP-43^WT^, hsa. TDP-43^G294V^ or hsa. TDP-43^dNLS^ transgenic lines (normalised to GAPDH, n = 3 replicates, p 2 0.5). **E.** Overexpression of hsa.TDP-43 did not significantly downregulate endogenous zebrafish TDP-43 (zZTDP-43) (normalised to GAPDH, n = 3 replicates, p 2 0.6). **F.** The hsa.TDP-43 transgene is expressed at a lower level than endogenous TDP-43 (hsa. TDP-43^WT^ mean = 38% of endogenous levels, hsa. TDP-43^G293V^ mean =59% of endogenous levels, and hsa. TDP-43^dNLS^ mean = 75% of endogenous levels (n = 3 replicates).

Neuronal overexpression of hsa.TDP-43^dNLS^ in a mouse model has been shown to downregulate endogenous mouse TDP-43 expression by approximately 50% due to TDP-43 autoregulation [33]. The hsa.TDP-43^dNLS^ mouse model therefore reflects concomitant cytoplasmic accumulation and nuclear depletion of TDP-43. To determine whether a similar downregulation occurs in zebrafish, we next quantified endogenous TDP-43 expression levels. Zebrafish carry two TDP-43 orthologues, zTDP-43 and zTDP-43-like (zTDP-43-L). Expression of both orthologues was quantified by western blot at 3 dpf, which showed no change in expression between non-transgenic controls (GAPDH normalised expression = 0.018), hsa.TDP-43^WT^ zebrafish (mean =0.031 ± 0.002), hsa.TDP-43^G294V^ zebrafish (mean = 0.023 ± 0.0008) or hsa.TDP-43^dNLS^ zebrafish (mean = 0.017 ± 0.0004) (**Fig 1 E**). Relative expression level of hsa.TDP-43 to endogenous TDP-43 demonstrated that the hsa.TDP-43 transgene is expressed at 38-75% of endogenous levels (**Fig 1F**).

To look for subtle changes in endogenous TDP-43 expression specifically in the central nervous system, western blot analysis was also performed on protein lysates collected from both brain and spinal cord of hsa.TDP-43 zebrafish at 3 months of age. No difference in expression level of either zTDP-43 orthologue was evident between non transgenic and hsa.TDP-43 samples (**Supplementary Fig 1**). These results indicate that, unlike the hsa.TDP-43^dNLS^ murine model, hsa.TDP-43^dNLS^ zebrafish display cytoplasmic accumulation without downregulation of endogenous (predominantly nuclear) zTDP-43 expression.

### hsa.TDP-43 expression did not significantly alter development of motor neurons or motor function at the embryonic stage

Zebrafish develop a functional motor system by 36 hpf [59]. We first investigated whether overexpression of hsa.TDP-43 affected this development. To facilitate this, transgenic zebrafish that express blue fluorescent protein (BFP) throughout the motor neuron soma and axons (*Tg*(*-3mnx1:tagBFP*)) [44] were crossed to the hsa.TDP-43 transgenic lines to enable visualisation of primary motor axon morphology (**Fig 2A**).

**Figure 2:**
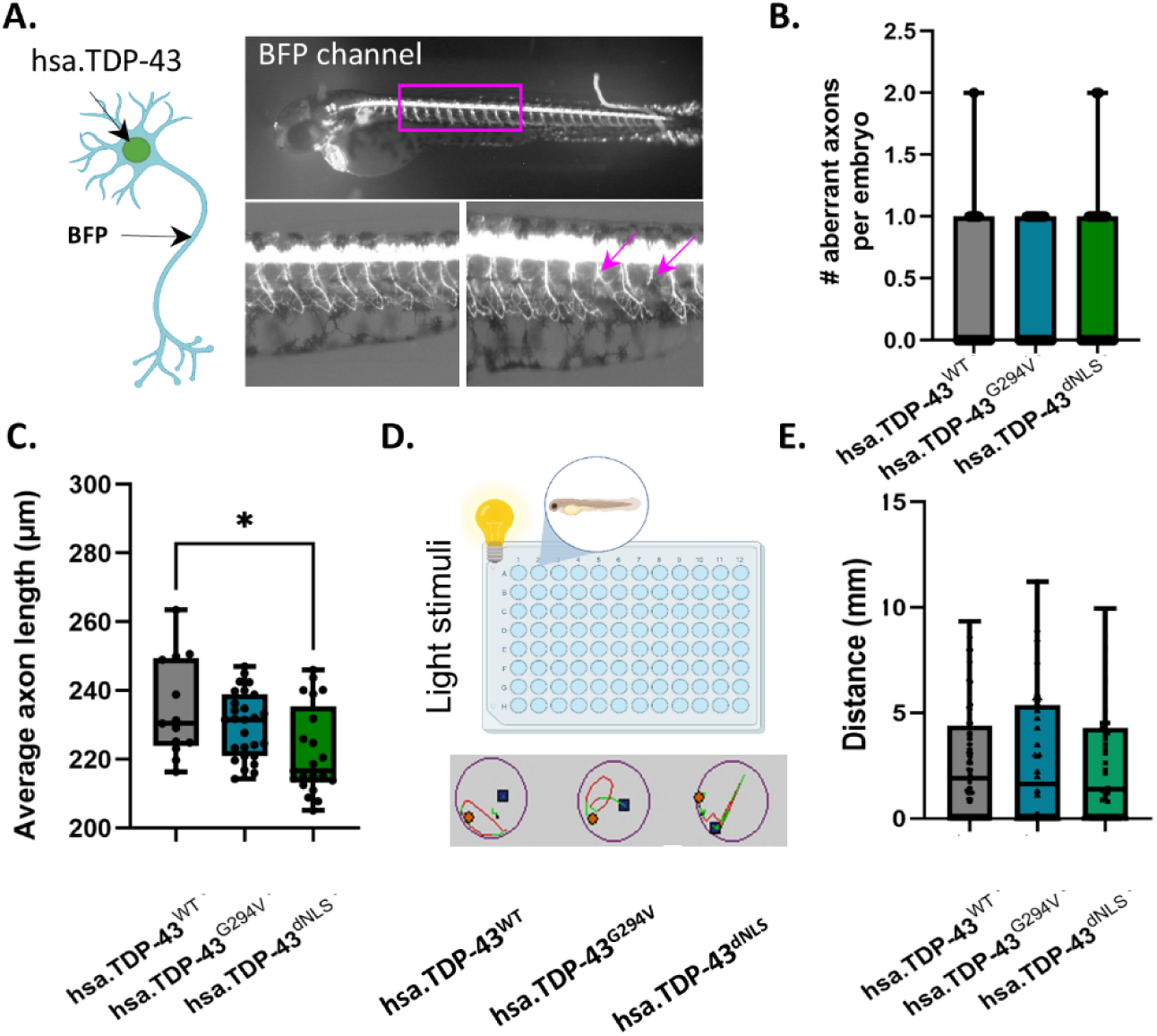
Characterisation of hsa.TDP-43 expression on motor neuron development and motor function. **A.** Schematic and representative images of hsa. TDP-43 transgenic fish at 2 days post fertilisation (dpf) that co-express blue fluorescent protein (BFP) throughout motor neuron soma and axon, as well as EGFP-tagged hsa. TDP-43. Bottom panel shows examples of primary motor axons with normal morphology (left panel) and axons with aberrant branching and truncation (arrows). **B.** Primary axon morphology was quantified at 2 dpf. No significant increase in incidence of aberrant morphology was observed between hsa. TDP-43^WT^ (mean = 0.32 +/- 0.11), hsa. TDP-43^G294V^ (mean = 0.37 +/- 0.09) or hsa. TDP-43^dNLS^ (mean = 0.48 +/- 0.12) zebrafish (n = 25-30 fish per group, Kruskal-Wallis analysis, p = 0.6). **C.** At2 dpf, a small, but significant reduction in axonal outgrowth was observed in hsa. TDP-43dNL$ zebrafish (mean = 222μm +/- 2.8) compared to hsa. TDP-43^WT^ zebrafish (mean = 235μm +/-4, p=0.01). No difference was observed in hsa. TDP-43^G294V^ zebrafish (mean = 230μm +/- 1.8) (n = 25-30 per group, one-way ANOVA, p = 0.48,). **D.** Schematic showing the method used to assess motor function at 2 dpf and representative traces. E. No difference in motor response to light stimulus was evident at 2 dpf between hsa. TDP-43^WT^ (mean = 2.6mm +/-0.27), hsa. TDP-43^G294V^ (mean = 2.8mm +/-0.4), and hsa. TDP-43^dNLS^ transgenic zebrafish (mean = 2.5mm +/-0.42), (n = 46-65 per group, Kruskal-Wallis test, p = 0.8).

Primary motor axon morphology at 48 hpf was examined for the presence of aberrant branching or truncation, which are established indicators of developmental defects [49] that correlate with impaired motor function [54], [60]. No differences in the incidence of aberrant primary axon morphology were observed between hsa.TDP-43^WT^ zebrafish (mean = 0.32 axons per fish ± 0.11), hsa.TDP-43^G294V^ zebrafish (mean = 0.37 ± 0.09) or hsa.TDP-43^dNLS^ zebrafish (mean = 0.48 ± 0.12) (**Fig 2B**). A mild, but statistically significant, difference in average length of primary axons was observed between the hsa.TDP-43^WT^ zebrafish (mean length = 229 μm ± 3) and hsa.TDP-43^dNLS^ zebrafish (mean length = 219 μm ± 2, p = 0.01) (**Fig 2C**). However, no difference in length was observed between hsa.TDP-43^WT^ or hsa.TDP-43^G294V^ zebrafish (mean length = 224 μm ± 2) (**Fig 2C**).

Next, we assessed motor function at 48 hpf. Spontaneous motor function is minimal at this timepoint [59], so the distance swum by embryos was assessed in response to a light stimulus (photomotor response) as previously described [54] (**Fig 2D**). No significant difference in distance swum in response to the stimulus was observed between hsa.TDP-43^WT^ zebrafish (mean = 2.6 mm ± 0.27), hsa.TDP-43^G294V^ (mean = 2.8 mm ± 0.4), and hsa.TDP-43^dNLS^ transgenic zebrafish (mean = 2.5 mm ± 0.42) (**Fig 2E**).

Collectively, these results indicate that while there is a mild defect in axon outgrowth in the hsa.TDP-43^dNLS^ zebrafish, all hsa.TDP-43 zebrafish showed normal morphological development and normal motor function at 2 dpf. Further, birefringence analysis of myotome integrity showed no defects in myofibrillar alignment at 6 dpf, indicating no effect of hsa.TDP-43 expression on muscle development (**Supplementary Fig 2**).

### Axonal transport speed was reduced in hsa.TDP-43^G294V^ zebrafish

Axonal transport deficits are reported to be an early feature of ALS, preceding development of gross motor neuron abnormalities and neuronal death [61]. To investigate whether defects are an early feature of our hsa.TDP-43 zebrafish, we next generated constructs to express mCherry-tagged KIF5A in motor neurons under the pan-neuronal *elavl3* promoter. KIF5A is a motor protein responsible for active axonal transport of organelles, RNA and proteins [62], thus providing a marker of general axonal transport.

The KIF5A construct was injected into hsa.TDP-43 eggs at the single cell stage of development, resulting in mosaic expression of mCherry-KIF5A in hsa.TDP-43 zebrafish. Live imaging of zebrafish positive for both hsa.TDP-43 and KIF5A was performed at 3 dpf (**Fig 3A**). The movement speed of KIF5A positive granules (0.8 – 2 µm^2^ in size) was tracked using ImageJ MTrackJ (**Fig 3B**). The average rate of transport of KIF5A particles in non-transgenic zebrafish axons was 1.6 µm/s ± 0.1 [62], [63]. No difference in average speed of KIF5A granules was evident between these non-transgenic controls and hsa.TDP-43^WT^ axons (1.4 µm/s ± 0.2) or hsa.TDP-43^dNLS^ axons (1.4 µm/s ± 0.41). However, a significant reduction in average speed was observed in axons of hsa.TDP-43^G294V^ zebrafish (0.96 µm/s ± 0.26, p <0.0001) (**Fig 3C**).

**Figure 3:**
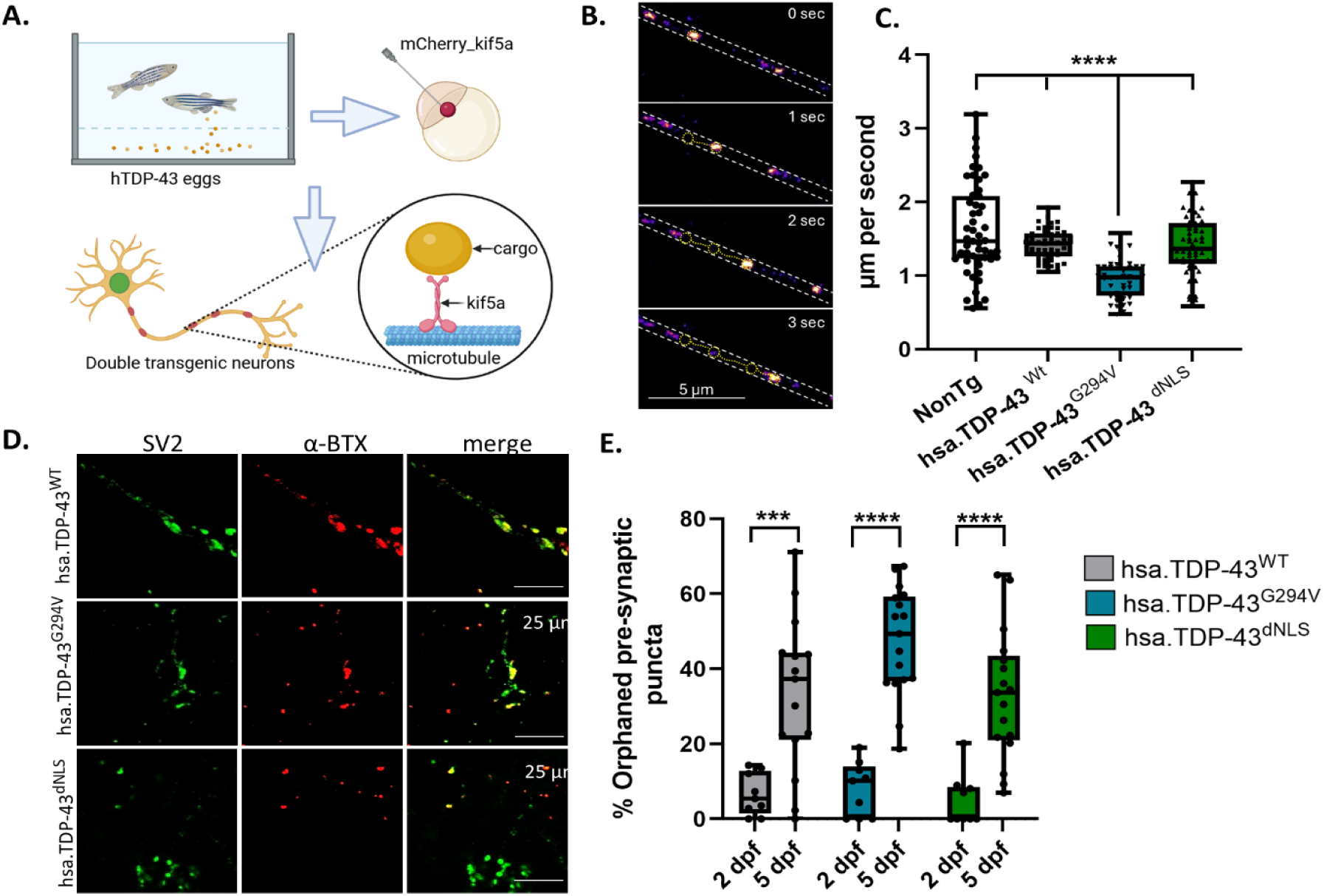
hsa.TDP-43 zebrafish show impaired axonal transport and neuromuscular junction. **A.** Schematic of method used to assess axonal transport in hsa. TDP-43 zebrafish. DNA encoding the motor protein kif5a tagged with mCherry was injected into eggs of hsa. TDP-43 zebrafish to produce double transgenic neurons. Movement of mCherry-kif5a along motor neuron axons was then quantified at 3 dpf. **B.** Representative images of timelapse series showing kif5a movement over 3 seconds. **C.** Kif5a movement was significantly slower (p < 0.0001) in hsa.TDP-43^G294V^ zebrafish (mean = 0.96 um/s = 0.26, n = 45) compared to non-transgenic controls (1.6 pm/s £ 0.1, n = 51), hsa. TDP-43^WT^ zebrafish (mean = 1.4 ym/s = 0.2, n = 57) and hsa.TDP-43^dNLS^ zeprafish (mean = 1.4 pm/s = 0.41, n = 51). The difference between non-transgenic controls and hsa.TDP-43^dNLS^ zebrafish approached, but did not reach significance (p = 0.058) (One-way ANOVA). D-E. Neuromuscular junction (NMJ) integrity was assessed at 2 dpf and 5 dpf by immunostaining with SV2 (pre-synaptic terminal) and a-bungarotoxin (a-BTX, post synaptic terminal). **D.** Representative images of NMJ staining in 5 dpf zebrafish shown. E. At 2 dpf, no difference in percentage of orphaned pre-synaptic puncta was evident between hsa. TDP-43^WT^ (mean = 6.6%, +/- 1.9) hsa. TDP-43^G294V^ (mean = 8.1%, +/-2.4) or hsa.TDP-43^dNLS^ (mean = 4.9% +/-2.3) (n = 9-10 embryos per group, One-way ANOVA, p = 0.6). By 5 dpf, the percentage of orphaned pre-synaptic puncta had significantly increased for all transgenic lines, with hsa. TDP-43^G294V^ showing the greatest increase (mean = 47% +/- 3.5) compared to hsa. TDP-43^WT^ (mean =33%, +/- 5.3) and hsa.TDP-43dNLS (mean = 33% +/- 4.2).

### Defects at the neuromuscular junction developed in hsa.TDP-43^G294V^ zebrafish

Defects in the NMJ have been reported to precede motor neuron loss in ALS [64] and cognitive impairment in AD [65]. We therefore examined the integrity of the NMJ in the hsa.TDP-43 transgenic lines. For this analysis, embryos were fixed at either 2 dpf or 5 dpf, and the presynaptic membrane was immunostained with an SV2 antibody and the postsynaptic membrane labelled with α-bungarotoxin (representative images at 5 dpf shown in **Fig 3D**). To identify disruption to the NMJ, the percentage of pre-synaptic puncta (nerve terminals) that did not have co-localised postsynaptic receptors, termed orphaned pre-synaptic puncta, was quantified.

At 2 dpf, the percentage of orphaned pre-synaptic puncta was similar across all hsa.TDP-43 transgenic lines (hsa.TDP-43^WT^ mean = 6.6% ± 1.9, hsa.TDP-43^G294V^ mean = 8.1% ± 2.4, hsa.TDP-43^dNLS^ mean = 4.9% ± 2.3) (**Fig 3E**). By 5 dpf, the number of orphaned pre-synaptic puncta had significantly increased for all transgenic lines, with hsa.TDP-43^G294V^ showing the greatest increase (hsa.TDP-43^WT^ mean = 33% ± 5.3, hsa.TDP-43^G294V^ mean = 47% ± 3.5, hsa.TDP-43^dNLS^ mean = 33% ± 4.2). The difference between hsa.TDP-43^WT^ and hsa.TDP-43^G294V^ was the most pronounced (p = 0.051, one-way ANOVA, **Fig 3E**). These changes in the pre-synapse/post-synapse ratio were not driven by a unilateral alteration of the pre- or post-synapse (**Supplementary Fig 3**).

### Cytoplasmic concentration of hsa.TDP-43 progressively increased in hsa.TDP-43^WT^ and hsa.TDP-43^G294V^ zebrafish

We previously demonstrated no difference in the TDP-43 nuclear-cytoplasmic ratio (N/C ratio) in hsa.TDP-43^WT^ and hsa.TDP-43^G294V^ zebrafish at 3 dpf^38^. This finding was replicated in the current study (hsa.TDP-43^WT^ mean = 3.6 ± 0.08 and hsa.TDP-43^G294V^ mean = 3.8 ± 0.086, p > 0.99). We next examined whether the hsa.TDP-43^WT^ and hsa.TDP-43^G294V^ zebrafish showed a progressive cytoplasmic accumulation of TDP-43 in motor neurons over time (‘early ageing’).

At 6 dpf, again no difference in the nuclear-cytoplasmic ratio was observed between hsa.TDP-43^WT^ and hsa.TDP-43^G294V^ (mean = 2.8 ± 0.06 and mean = 2.7 ± 0.08 respectively, p>0.99). However, the absolute nuclear-cytoplasmic ratios were significantly reduced between timepoints for both hsa.TDP-43^WT^ (p < 0.0001) and hsa.TDP-43^G294V^ (p =0.0006, **Fig 4B**). Examination of compartment-specific expression of TDP-43 demonstrated that the relative nuclear expression reduced between 3 dpf and 6 dpf (1.86 ± 0.03 to 1.6 ± 0.03), while relative cytoplasmic TDP-43 increased (0.497 ± 0.008 to 0.557 ± 0.007, **Supplementary Fig 4**). Collectively, these results indicate a progressive cytoplasmic shift of TDP-43 over time – leaving the nucleus and accumulating in the cytoplasm.

**Figure 4:**
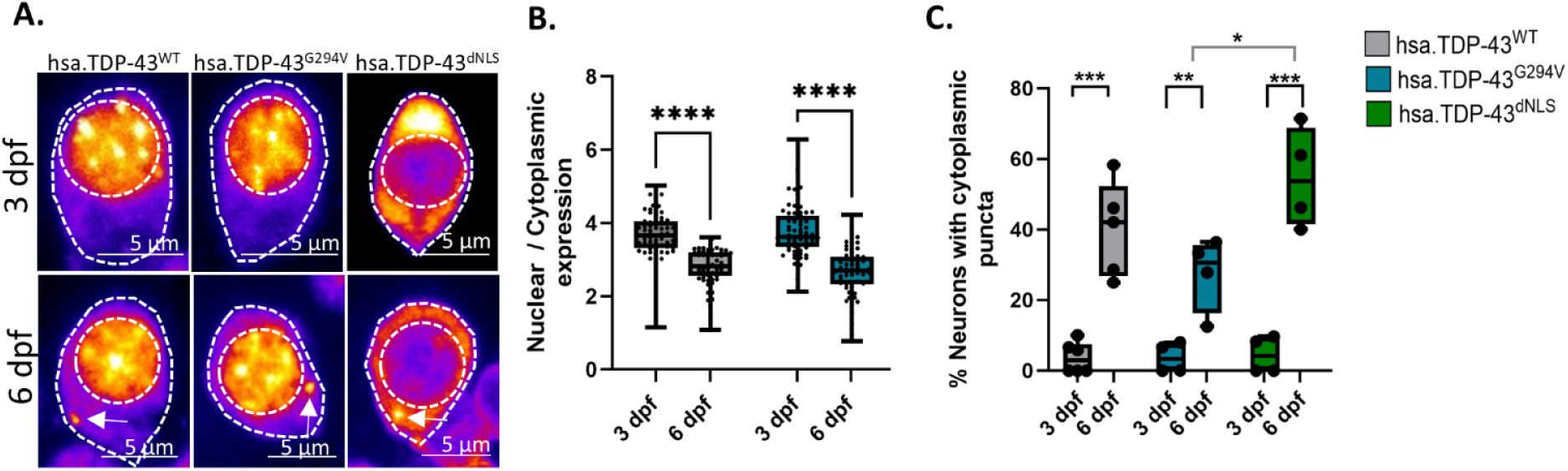
Nuclear hsa.TDP-43 progressively shifts to the cytoplasm and cytoplasmic hsa.TDP-43 forms discrete puncta. **A.** Representative images showing nuclear and cytoplasmic expression of TDP-43 variants and examples of cytoplasmic puncta that develop by 6 dpf. **B.** Nuclear-cytoplasmic ratio of hsa. TDP-43 was quantified at 3 dpf and 6 dpf in the hsa. TDP-43^WT^ and hsa. TDP-43^G294V^ zebrafish. No significant difference in localisation between hsa. TDP-43^WT^ and hsa. TDP-43^G294V^ was observed at either 2 dpf (mean = 3.6 = 0.08 and mean = 3.8 = 0.086 respectively, p > 0.99) or 6 dpf (mean = 2.8 + 0.06 and mean = 2.7 = 0.08 respectively, p > 0.99). However, a significant cytoplasmic shift was evident from 3 dpf to 6 dpf in both hsa. TDP-43^WT^ (p < 0.0001) and hsa. TDP-43^G294V^ (p= 0.0006) zebrafish (n = 59-71 per group, Kruskal-Wallis analysis). **C.** Quantification of the percentage of primary motor neuron soma per fish that contained cytoplasmic puncta. The number of puncta significantly increased between 3 dpf and 6 dpf in all hsa.TDP-43 lines (hsa.TDP-43^WT^3 dpf mean = 3.7% +/-1.8, 6 dpf mean = 40% +/- 6, hsa. TDP-43^G294V^ 3 dpf mean = 3.7% +/- 2.1, 6 dpf mean = 28% +/- 5.3 and hsa. TDP-43^dNLS^ 3 dpf mean = 4.5% +/- 2.6, 6 dpf mean 55% +/- 7.1, unpaired t-tests). hsa. TDP-43^dNLS^ had significantly more puncta than hsa. TDP-43^G294V^ zebrafish at 6 dpf (p = 0.035, Oneway ANOVA) (n = 5 fish per group, 51-73 neurons per fish).

Next, we assessed the formation of cytoplasmic hsa.TDP-43 puncta (conceivably precursors of aggregates) in zebrafish motor neurons over this timeframe. Cytoplasmic puncta were rarely observed at 3 dpf (hsa.TDP-43^WT^ average neurons with puncta = 3.7 % ± 1.8, hsa.TDP-43^G294V^ average neurons with puncta = 3.7 % ± 2.1, hsa.TDP-43^dNLS^ average neurons with puncta = 4.6 % ± 2.1). However, their frequency significantly increased in all hsa.TDP-43 transgenic lines by 6 dpf (**Fig 4C**). Cytoplasmic puncta were most commonly observed in hsa.TDP-43^dNLS^ (average number of neurons with puncta = 55 % ± 7.1), followed by hsa.TDP-43^WT^ zebrafish (average number of neurons with puncta = 40 % ± 6 and then hsa.TDP-43^G294V^ zebrafish (average number of neurons with puncta = 28 % ± 5.3) (**Fig 4C**).

Overall, the progressive increase in cytoplasmic hsa.TDP-43 and formation of discrete puncta indicates that the hsa.TDP-43 models may prove useful tools for longitudinal analysis of the development of hallmark TDP-43 pathologies such as liquid-to-solid transition, cytoplasmic sequestration, and aggregation.

### hsa.TDP-43 zebrafish showed reduced motor function at 6 days post fertilisation

By 6 dpf, zebrafish larvae exhibit substantive spontaneous movement and a variety of evoked motor behaviours. Exposure to repeated light-dark cycles has been shown to elicit an anxiety-like response resulting in increased activity, thereby providing a reliable readout of motor function and fatigue [55], [66]. To assess the hsa.TDP-43 zebrafish, the distance swum at 6 dpf was recorded over a 20-minute period consisting of 5 minute light-dark cycles (**Fig 5A**). The hsa.TDP-43^WT^ zebrafish swam significantly greater distances (mean = 2613 mm ± 82) than both hsa.TDP-43^G294V^ zebrafish (mean = 2292 mm ± 63, p = 0.0016) and hsa.TDP-43^dNLS^ zebrafish (mean = 2470 mm ± 66, p = 0.0001) (**Fig 5B**). Non-transgenic zebrafish swum significantly further than all hsa.TDP-43 groups (mean = 3451 mm ± 69, p ≤ 0.0001) (**Fig 5B**).

**Figure 5:**
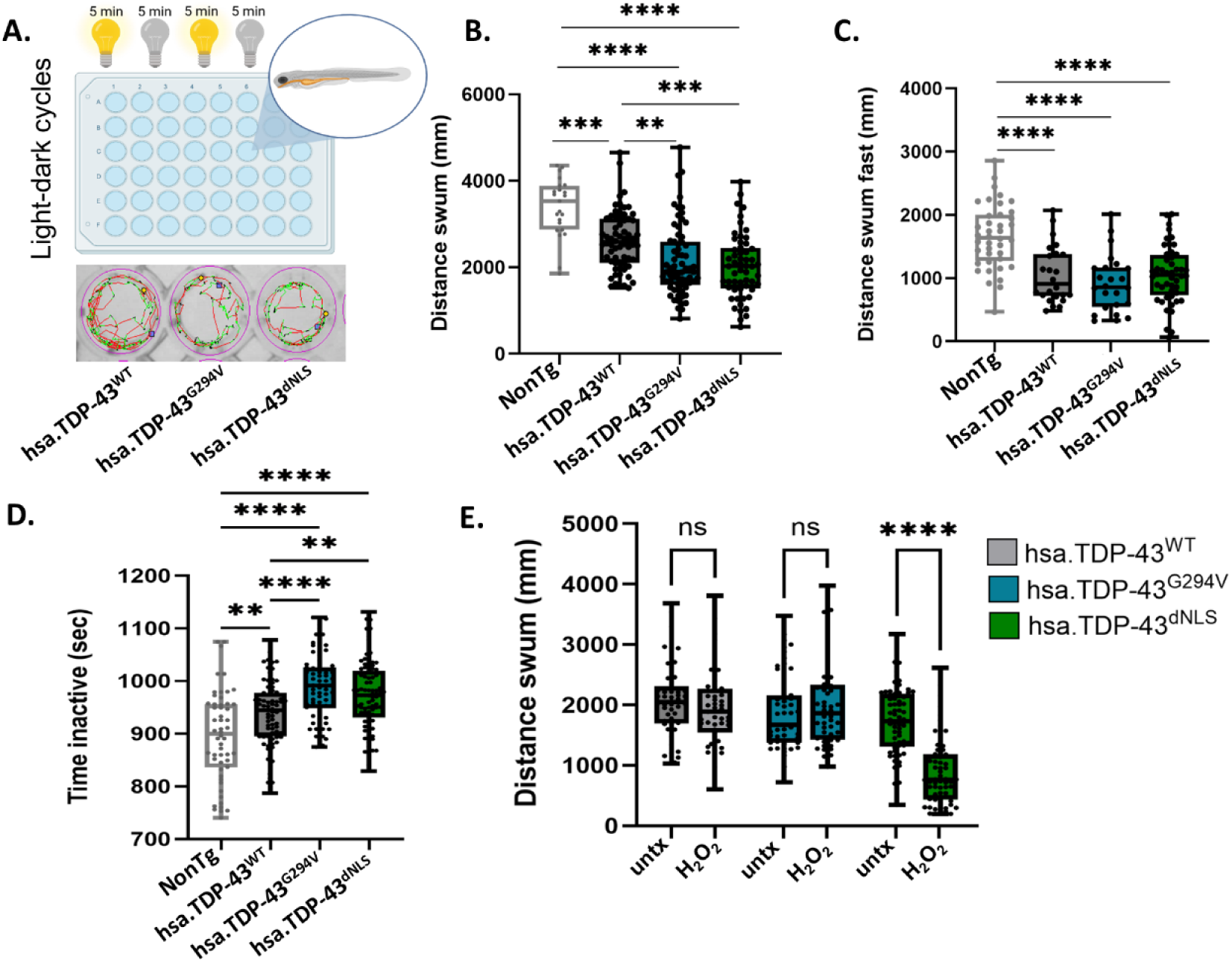
hsa.TDP-43 zebrafish show reduced motor function at 6 days post fertilisation. **A.** Schematic showing method used to assess motor function at 6 dpf and representative traces. **B.** All hsa. TDP-43 transgenic lines swum significantly shorter distances than non-transgenic controls (mean = 3451mm +/- 69). Both hsa. TDP-43^dNLS^ (mean = 2470mm +/- 66) and hsa. TDP-43^G294V^ zeprafish (mean = 2292mm +/- 63) swum shorter distances than hsa. TDP-43^WT^ zebrafish (mean = 2788mm +/- 69) (n = 56-66 per group, Kruskal-Wallis test). **C.** Non-transgenic zebrafish swum significantly greater distances at high speed compared to all hsa. TDP-43 groups. (mean = 1658mm +/- 81). No difference in distance swum at high speed was observed between hsa. TDP-43^WT^ zebrafish (mean = 1056 +/- 92), hsa. TDP-43^G294V^ zebrafish (mean = 916 +/- 92) or hsa. TDP-43^dNLS^ (mean = 1051 +/- 65) (n = 46—65 per group, One-way ANOVA, p = 0.44). **D.** All hsa.TDP-43 zebrafish spent significantly more time at rest than non-transgenic zebrafish (mean = 895 sec +/- 12, p = 0.001). Both hsa.TDP-43^G294V^ zebrafish (mean = 990 sec +/- 7.5) and hsa. TDP-43^dNLS^ (mean = 972 sec +/- 7.1) spent significantly more time at rest compared to hsa. TDP-43^WT^ zebrafish (mean = 937 sec +/- 7.1, p = 0.0006 and p = 0.066 respectively). **E.** Reduced motor function was exacerbated by oxidative stress in hsa. TDP-43^dNLS^ zebrafish (mean untreated = 1723mm +/- 69, H,0, treated = 897 mm +/- 74). The effect of oxidative stress in hsa. TDP-43^WT^ zebrafish (mean untreated = 2076 mm +/- 88, H_2_,0_2_, treated = 1943 mm +/- 90) and hsa. TDP-43^G294V^ zebrafish (untreated mean = 1890 mm +/- 95, H,0, treated = 1999 mm +/- 103) did not reach significance (p =0.3 and p = 0.17 respectively).

To determine whether the reduced distance swum was a consequence of a reduced ability to swim quickly or due to increased fatigue, distance swum at high speed (>10 mm / minute) and time spent at rest were examined. Non-transgenic zebrafish were found to swim significantly further distances at top speed compared to all hsa.TDP-43 zebrafish (mean = 1658 mm ± 81, p < 0.0001). However, no differences in distance swum at high speed were observed between hsa.TDP-43^WT^ zebrafish (mean = 1056 mm ± 92), hsa.TDP-43^G294V^ zebrafish (mean = 916 mm ± 93) or hsa.TDP-43^dNLS^ zebrafish (mean = 1051 mm ± 65) (**Fig 5C**). Non-transgenic zebrafish also spent less time at rest compared to all hsa.TDP-43 zebrafish (mean = 895 sec ± 12, p = 0.001). However, the amount of time spent at rest was significantly higher in both the hsa.TDP-43^G294V^ zebrafish (mean = 990 sec ± 7.5) and hsa.TDP-43^dNLS^ (972 sec ± 7.1) zebrafish compared to hsa.TDP-43^WT^ zebrafish (mean = 943 sec ± 7.8, p =0.0006 and p = 0.066 respectively) (**Fig 5D**).

hsa.TDP-43 variants was sufficient to affect motor function as indicated by reduced swimming speed and increased fatigue. These phenotypes were more pronounced with cytoplasmic accumulation of hsa.TDP-43 or nuclear expression of an ALS-causative mutation compared to nuclear overexpression of wildtype hsa.TDP-43.

### Motor dysfunction was exacerbated by oxidative stress in hsa.TDP-43^dNLS^ zebrafish

Oxidative stress is a common feature of neurodegenerative diseases and has been shown to promote TDP-43 mislocalisation, hyperphosphorylation and aggregation [67]–[69], resulting in impaired RNA binding ability [67] and mitochondrial dysfunction [69]. To examine the sensitivity of the hsa.TDP-43 zebrafish to oxidative stress, each transgenic line was exposed to 0.5 mM hydrogen peroxide from 3 dpf (refreshed daily) and motor function assessed at 6 dpf (**Fig 5E**).

Notably, hsa.TDP-43^dNLS^ zebrafish demonstrated a marked sensitivity to oxidative stress as indicated by a significant reduction in distance swum in the treated group compared to their untreated clutch-mates (untreated mean = 1723 mm ± 69, H_2_O_2_ treated mean = 897 mm ± 74, p < 0.0001; **Fig 5E**). In comparison, hsa.TDP-43^WT^ zebrafish (untreated mean = 2076 mm ± 88, H_2_O_2_ treated mean = 1943 mm ± 90) and hsa.TDP-43^G294V^ zebrafish (untreated mean = 1890 mm ± 95, H_2_O_2_ treated mean = 1999 mm ± 103) did not show a reduced motor function after exposure to oxidative stress (p = 0.99 and p > 0.99 respectively) at 6 dpf.

Cellular characterisation of hsa.TDP-43 expression in primary motor neurons of zebrafish treated with H_2_O_2_ did not reveal significant differences with respect to nuclear-cytoplasmic ratio or puncta number between treated and untreated clutch mates in any of the hsa.TDP-43 zebrafish lines (**Supplementary Fig 5**).

### Effect of ALS-approved drugs on the hsa.TDP-43 zebrafish motor phenotype

Riluzole and edaravone are two FDA approved drugs for the treatment of ALS – riluzole, an anti-glutamatergic drug targets neuronal hyperexcitability [70], and edaravone is known to have potent antioxidant effects [71].

#### Riluzole had an adverse effect on hsa.TDP-43^WT^ motor function

In order to examine whether neuronal hyperexcitability is a driver for motor impairment in the hsa.TDP-43 zebrafish, we tested the effect of riluzole on motor function at 6 dpf. Larvae were dechorionated at 24 hpf, clutches split into two and water treated with either riluzole (7.5 µM) or DMSO as previously described [57]. At 6 dpf, motor function was assessed using the 20 minute light-dark protocol. No statistically significant difference in distance swum was observed between DMSO and riluzole treated zebrafish from either the hsa.TDP-43^G294V^ (DMSO mean = 2039 mm ± 84, riluzole mean = 2125 mm ±74) or hsa.TDP-43^dNLS^ zebrafish (DMSO mean = 2023mm ± 99, riluzole mean = 1999 ±87). However, riluzole exacerbated the swimming phenotype in hsa.TDP-43^WT^ zebrafish by significantly reducing the distance swum (DMSO mean = 2553 ± 73, riluzole mean = 2250 ± 71, two-tailed t-test, p = 0.001) (**Fig 6A**).

**Figure 6:**
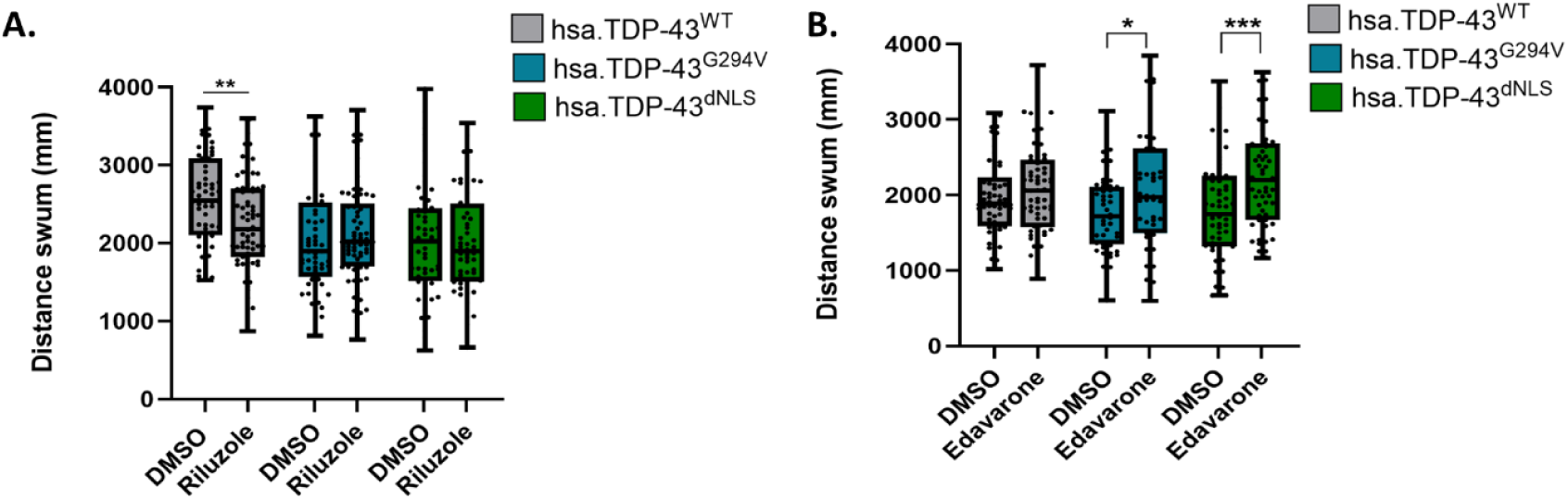
Riluzole does not improve motor function of hsa.TDP-43 zebrafish but Edaravone does. **A.** Treatment with 7.5 pM Riluzole did not alter the distance swum in hsa. TDP-43^G294V^ zebrafish (DMSO mean = 2039 mm = 84, Riluzole mean = 2125 mm = 74, p >0.99, n = 68-69) or hsa. TDP-43^dNLS^ zebrafish (DMSO mean = 2023 mm * 99, Riluzole mean =1999 mm # 87 p >0.99, n = 56). However, Riluzole treatment did reduce the distance swum in hsa.TDP-43^WT^ fish (DMSO mean = 2553 + 73, Riluzole mean = 2250 = 71, p = 0.001, n = 64-66, two-tailed t-test). **B.** Treatment with 5 uM Edaravone was associated with a significant rescue in distance swum in the hsa. TDP-43^dNLS^ zebrafish (DMSO mean = 1794 mm +/- 84, Edaravone mean = 2241 mm +/- 88, p = 0.0003, unpaired t-test, n = 59) and to a lesser extent, hsa. TDP-43^G294V^ zebrafish (DMSO mean =1755 mm +/- 71 Edaravone mean =2084 mm +/-119, p = 0.019, unpaired t-test, n = 47-48) but not in hsa. TDP-43^WT^ (DMSO mean =1975 mm +/- 62, Edaravone mean =2082 mm +/- 75, p = 0.3, Mann-Whitney test, n = 68).

#### Edaravone improved motor function of hsa.TDP-43^dNLS^ zebrafish

Edaravone has been reported to improve motor function in a toxin-induced zebrafish model at a single dose of 100 µM applied at 4 hpf [72]. However, to establish an appropriate dose of edaravone for daily treatment from 3 dpf in dechorionated embryos, a dose-response trial was performed in hsa.TDP-43^dNLS^ zebrafish. Edaravone at 5–100 µM was added to the water of 3 dpf embryos and refreshed daily. Doses > 40 µM were found to be toxic and 5 µM edaravone was identified as sufficient to affect motor function (**Supplementary Fig 6**).

Following treatment with edaravone (5 µM) or DMSO from 3 dpf, motor function was assessed at 6 dpf using the 20 minute light-dark protocol. Edaravone treated hsa.TDP-43^dNLS^ zebrafish showed a statistically significant increase in distance swum compared to clutch-mate DMSO treated controls (DMSO mean = 1794 ± 84, edaravone mean = 2241 ± 88, p = 0.0003). Hsa.TDP-43^G294V^ zebrafish also showed a significant recovery in distance swum with edaravone treatment (DMSO mean = 1755 +/- 71, edaravone mean = 2084 ± 119, p = 0.019). No rescue of the swimming effect was evident in edaravone-treated hsa.TDP-43^WT^ zebrafish (DMSO mean = 1975 ± 62, edaravone mean = 2082 ± 75, p = 0.3) (**Fig 6B**).

### Aged hsa.TDP-43 zebrafish showed reduced lifespan and loss of motor neurons

#### Survival analysis

To investigate the long-term effects of hsa.TDP-43 expression, the F3 and F4 generations of hsa.TDP-43 transgenic zebrafish were raised to maturity and aged to 14 months. A significant reduction in survival was evident in both the hsa.TDP-43^dNLS^ and hsa.TDP-43^G294V^ zebrafish compared to the hsa.TDP-43^WT^ zebrafish (**Fig 7A**; p = 0.0001 and p = 0.0002 respectively, n = 74 per group, Log-rank test for comparison of survival curves).

**Figure 7:**
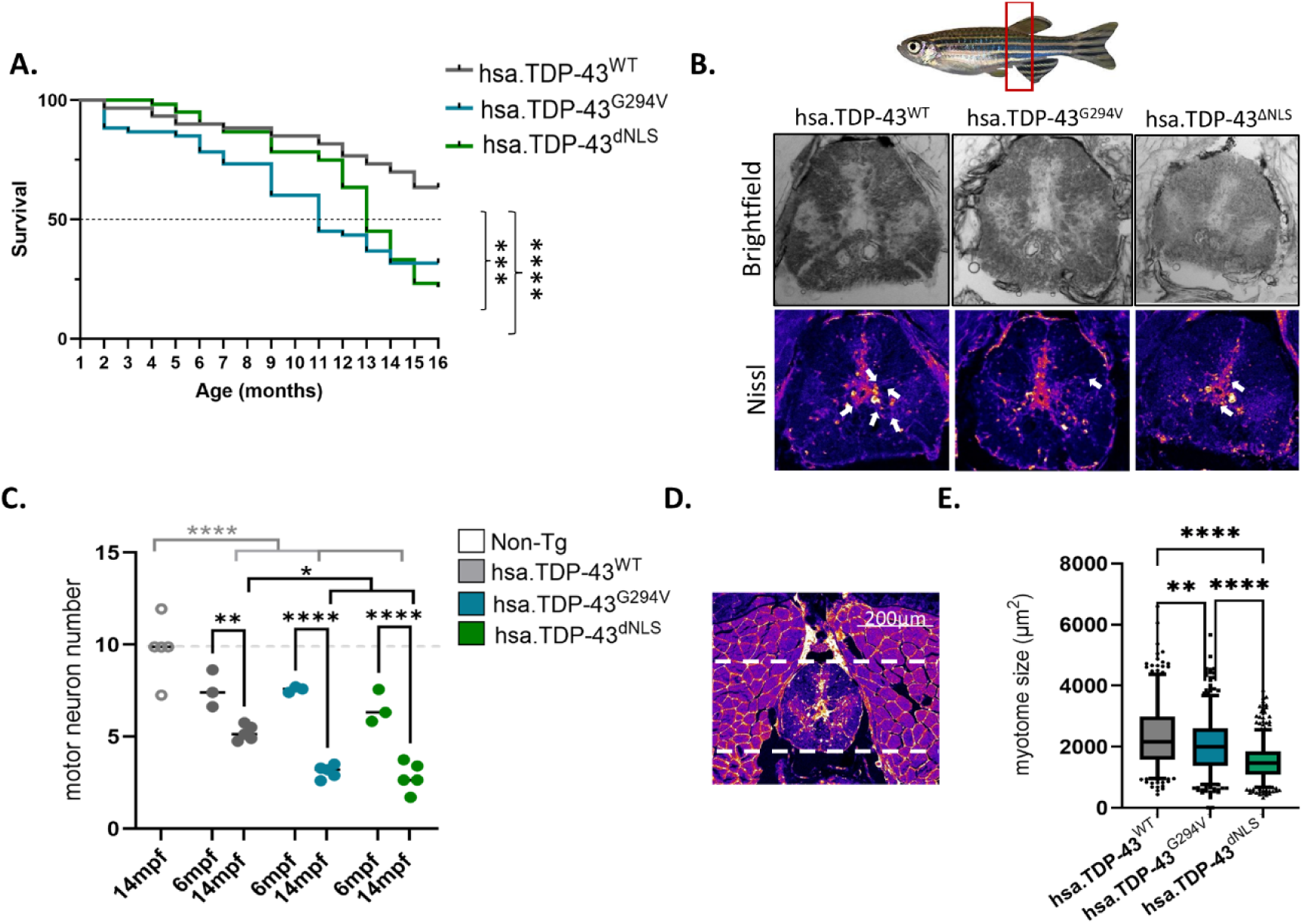
Overexpression of hsa.TDP-43 is associated with reduced survival and motor neuron loss. **A.** Reduced survival of hsa.TDP-43^dNLS^ zebrafish (median survival= 11 months) and hsa.TDP-43^G294v^ zebrafish (median survival = 13 months) relative to hsa.TDP-43^WT^ zebrafish was observed (p = 0.001 and p = 0002 respectively, Log-rank Mantel-Cox test, n = 74 per group). **B.** Representative images of Nissl-stained spinal cord cross sections used for motor neuron counts. **C.** Quantification of the number of large (>75 µm^2^) Nissl-stained neurons within the ventral horn was performed at 6 mpf and 14 mpf. A significant loss of motor neurons was observed between these timepoints in hsa.TDP-43^WT^ zebrafish (6 mpf mean= 7.6 ± 0.6, 14 mpf mean= 5.2 ± 0.2), hsa.TDP-43^G294v^ (6 mpf mean =7.6 ± 0.09, 14 mpf mean = 3.1 ± 0.2) and hsa.TDP-43^dNLS^ (6 mpf mean= 6.6 ± 0.5, 14 mpf mean =2.8 ±0.35 n = 3 fish, n >20 sections per zebrafish, 3-5 zebrafish One-way ANOVA). At 14 mpf, the number of large motor neurons was significantly lower in both hsa.TDP-43^dNLS^ (mean =2.8 ± 0.35) and hsa.TDP-43^G294v^ zebrafish (mean= 3.1 ± 0.16) compared to hsa.TDP-43^WT^ zebrafish (mean= 5.2 ± 0.2). Non-transgenic controls retained significantly more motor neurons than all hsa.TDP-43 transgenic zebrafish (mean = 9.8 ± 0.7, one-way ANOVA). **D.** Representative image showing epaxial muscles included in analysis of myotome size at 14 mpf. Area of muscle segments within the dorsal and ventral bounds of the spinal cord (dashed line) and within 200 µm of the spinal cord were quantified. **E.** Average myotome size was significantly reduced in both hsa.TDP-43^dNLS^ (mean= 1509 µm^2^ ± 55) and hsa.TDP-43^G294V^ (mean= 2056 µm^2^ ± 43) zebrafish compared to hsa.TDP-43^WT^ zebrafish (mean = 2355 µm^2^ ± 55, n = 3 sections per fish, 5 fish per group, Kruskal-Wallis test).

#### Motor neuron counts

To determine the effects of hsa.TDP-43 expression on motor neuron survival, transgenic zebrafish were euthanised at 6 and 14 mpf for histological examination (**Fig 7B**). Following fixation and dehydration, 30 μm thick cryostat prepared sections of the segment directly ventral to the dorsal fin were assessed for motor neuron counts. The number of mature motor neurons, classified as Nissl stained cells >75 μm^2^ within the spinal cord ventral horns, was quantified using ImageJ as previously described [58], [73]. A minimum of 20 sections per fish was examined, and the average was recorded (**Fig 7C, 7D**).

At six months of age, no significant difference in mean number of mature motor neurons was evident between groups (n = 3 fish per group, hsa.TDP-43^WT^ mean = 7.6 ± 0.58, hsa.TDP-43^G294V^ mean = 7.6 ± 0.09, hsa.TDP-43^dNLS^ mean = 6.6 ± 0.52). At 14 months of age, a significant loss of motor neurons was evident in all three groups, with the mean number of motor neurons per segment in hsa.TDP-43^WT^ zebrafish reduced to 5.2 ± 0.19 (p = 0.0003), in hsa.TDP-43^G294V^ zebrafish dropping to 2.8 ± 0.35 (p <0.0001), and in hsa.TDP-43^dNLS^ zebrafish dropping to 2.8 ± 0.35 (p < 0.0001) (**Fig 7C**). At 14 months of age, the number of mature motor neurons was also significantly reduced in both the hsa.TDP-43^dNLS^ zebrafish and hsa.TDP-43^G294V^ zebrafish (mean = 2.8 ± 0.35) compared to hsa.TDP-43^WT^ zebrafish (p = 0.019 and p = 0.0075 respectively) (n = 5 zebrafish per group, One-way ANOVA) (**Fig 7C**). To examine how much of this motor neuron loss was associated with aging rather than overexpression of hsa.TDP-43, motor neuron counts were compared to non-transgenic zebrafish at 14 mpf. Non-transgenic zebrafish had significantly higher number of mature motor neurons at 14 mpf (mean = 9.8 ± 0.7) compared to hsa.TDP-43^WT^ (p <0.0001), hsa.TDP-43^G294V^ zebrafish (p <0.0001) and hsa.TDP-43^dNLS^ zebrafish (p <0.0001) (**Fig 7C**), indicating a significant role of hsa.TDP-43 expression, independent of variant, in motor neuron loss in zebrafish. Overall this reduction reflects a ∼47-72% loss of motor neurons compared to non-transgenic controls at 14 mpf (47% for hsa.TDP-43^WT^; 68.4% for hsa.TDP-43^G294V^; 72% for hsa.TDP-43^dNLS^).

#### Myotome size assessment

To examine the effect of loss of spinal cord motor neurons on the musculature, average myotome size at 14 mpf was quantified in the epaxial muscles immediately adjacent to the spinal cord in the Nissl-stained cross sections (**Fig 7D**). In line with the motor neuron analysis, a significant reduction in average myotome size was evident in both the hsa.TDP-43^dNLS^ zebrafish (mean = 1509 μm^2^ ± 55) and hsa.TDP-43^G294V^ zebrafish (mean = 2056 μm^2^ ± 43) compared to hsa.TDP-43^Wt^ zebrafish (mean = 2355 μm^2^ ± 55, p = 0.0007 and p<0.0001 respectively) (**Fig 7E**).

### Accumulation of phosphorylated TDP-43 was a feature of hsa.TDP-43 zebrafish

The formation of hyper-phosphorylated TDP-43 aggregates is considered the pathological hallmark of ALS [2], [3], FTLD [3], LATE [4] and is present in up to 57% of AD patients [74]. To determine whether this characteristic pathology develops in the hsa.TDP-43 transgenic zebrafish, 6 mpf zebrafish sections were immunostained with an antibody against endogenous (native) TDP-43 (nTDP-43) and a phospho-TDP-43 (pTDP-43) antibody, together with NeuroTrace Nissl stain, and DAPI as a nuclear marker (**Fig 8A**). Nuclear pTDP was not detected in non-transgenic zebrafish neurons. Both TDP-43^WT^ and TDP-43^dNLS^ zebrafish demonstrated a mild but not statistically significant increase in nuclear pTDP-43 (p = 0.066 and *p* = 0.07 respectively). TDP-43^G294V^ zebrafish displayed a significant increase in nuclear pTDP-43 relative to non-Tg controls (14.38 ± 16.86, *p* <0.0001) (**Fig 8A,B**).

**Figure 8:**
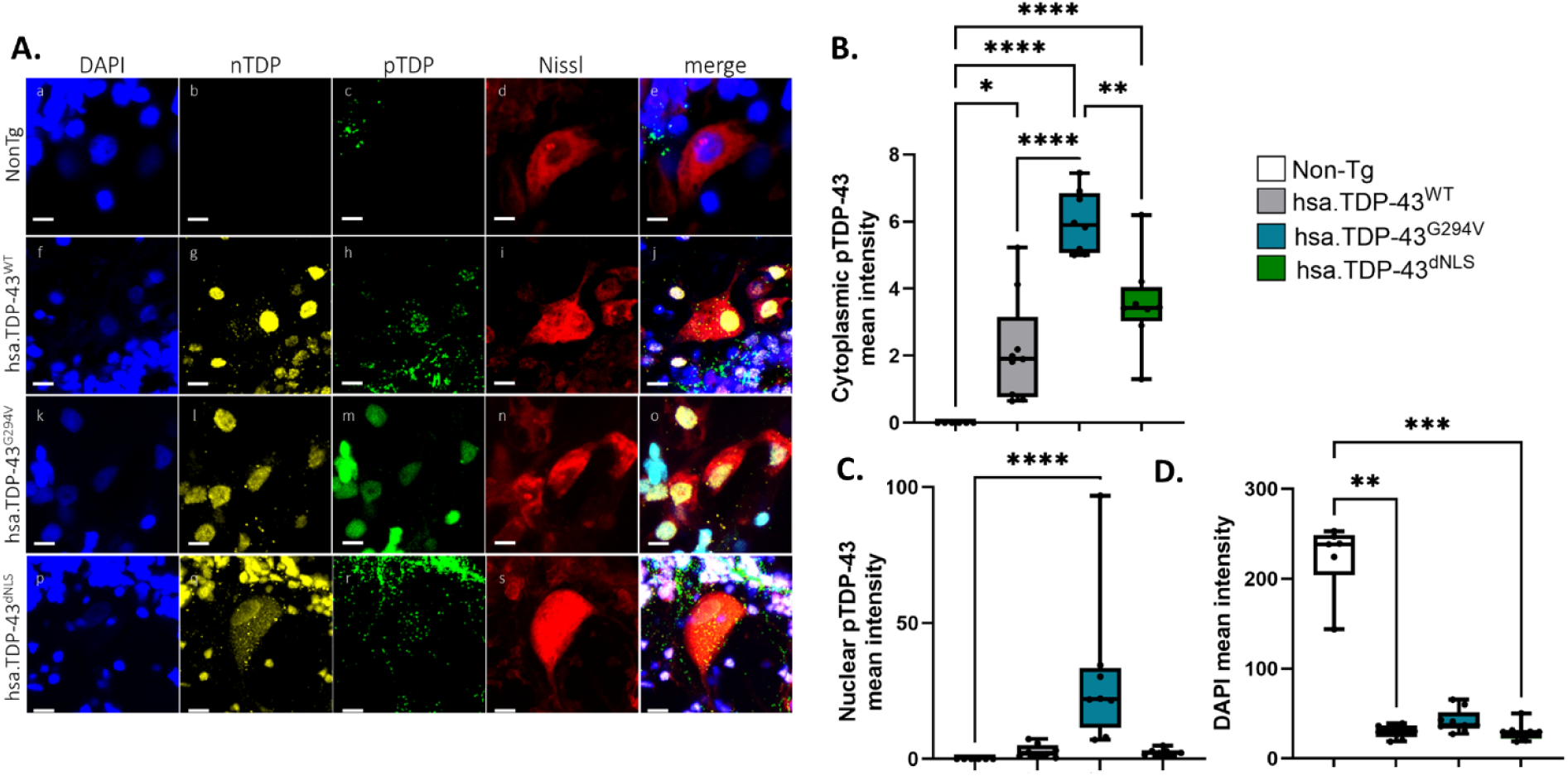
Aged hsa.TDP-43 zebrafish show neuronal accumulation of phospho-TDP-43. **A.** Representative images of adult (6 mpf) zebrafish spinal cord motor neurons immunostained with a native TDP-43 (nTDP-43) antibody which binds both hsa.TDP-43 and zTDP-43, and phospho-TDP-43 (pTDP-43) antibodt, nissl neuronal marker and DAPI nuclear marker. Scale bar = 10 pm. **B.** Nuclear pTDP was not detected in neurons of non-transgenic zebrafish (0.00 ± 0.00, n = 6). TDP-43^WT^ (2.9 ± 0.86 n = 9) and TDP-43^dNLS^ zebrafish (2.5 = 0.46, n = 9) showed a non-significant increase in nuclear pTDP-43 compared to non-transgenic controls (p = 0.066 and *p* = 0.07 respectively). A significant increase in nuclear pTDP-43 was observed in nuclei of TDP-43^G294V^ motor neurons (30 = 10, *p* <0001, n = 8, Kruskal-Wallis test). **C.** Cytoplasmic pTDP-43 expression was not detected in non-transgenic zebrafish (0.00 ± 0.00). However, all three hsa. TDP-43 transgenic lines showed significantly elevated cytoplasmic pTDP-43 expression (hsa. TDP-43^WT^ = 2.2±0.52, *p*=0.01), hsa.TDP-43^G294V^ (6 ± 0.33, p<0.0001) and hsa. TDP-43^dNLS^ = 3.6 ± 0.48, p<0.0001). pTDP-43 expression was significantly higher in hsa. TDP-43^G294V^ zebrafish compared to both hsa. TDP-43^WT^ (p<0.0001) and hsa. TDP-43^dNLS^ (p = 0.002, One-way ANOVA). **D.** The mean intensity of DAPI staining was quantified from the same images to gain insight into neuronal health. DAPI intensity was highest in the non-transgenic controls (224.3 ± 16.6). All hsa.TDP-43 zebrafish showed a significant reduction in intensity (hsa. TDP-43^WT^ = 29.97 = 2.4, p=0.0007, hsa.TDP-43^G294V^ = 42 04 ± 4.3, p =0.08 and hsa. TDP-43^dNLS^ = 29.8 ± 3.4, p =0.0004, Kruskal-Wallis test).

Cytoplasmic pTDP-43 expression was also not detected in non-transgenic zebrafish neurons. However, all three hsa.TDP-43 transgenic lines showed significantly elevated pTDP-43 expression in the cytoplasm (TDP-43^WT^ p = 0.01, TDP-43^G294V^ p <0.0001 and TDP-43^dNLS^ p<0.0001). (**Fig 8C**).

DAPI, as a DNA-binding stain, can provide insight into the DNA content, or lack thereof of cells. To investigate DNA content as a reflection of cell health [75], the mean intensity of DAPI staining was quantified. DAPI intensity was highest in the non-transgenic controls (220.6 ± 51.59). All hsa.TDP-43 zebrafish showed a significant reduction in DAPI staining intensity in comparison (TDP-43^WT^ = 27.2 ± 9.37, TDP-43^G294V^ =41.5 ± 16.63 and TDP-43^dNLS^ = 29.8 ± 14.59, p < 0.01). This indicated reduced intact DNA within hsa.TDP-43 neurons, which is consistent with our finding of reduced neuronal survival at this timepoint relative to non-transgenic controls.

## Discussion

We present a detailed assessment of stable transgenic zebrafish models that overexpress human TDP-43 (hsa.TDP-43) in neurons throughout the spinal cord. This characterisation revealed that motor neuron expression of hsa.TDP-43 induced distinct gain-of-function effects leading to ALS-relevant phenotypes, including reduced motor function by 6 dpf, susceptibility to oxidative stress, loss of motor neurons, muscle wastage, and reduced survival. The FDA-approved ALS drug Edaravone, but not Riluzole, was found to rescue motor impairment in young hsa.TDP-43^dNLS^ zebrafish. Collectively, these findings demonstrate that these neuron-specific zebrafish models provide a valuable tool for examining pathogenic mechanisms of TDP-43 pathology in neurodegenerative diseases and enabling efficient drug screening for therapeutic evaluation.

Zebrafish have been extensively used to model TDP-43 proteinopathies, with studies employing diverse approaches to examine different facets of TDP-43 pathology and elucidate its role in neurodegeneration. Ubiquitous knockdown of zebrafish TDP-43 has been shown to induce developmental defects and motor dysfunction [20], [21], studies that highlight the critical role of TDP-43 in RNA regulation and neuronal maintenance. Ubiquitous overexpression of an ALS-linked TDP-43 mutation (G348C) further highlighted the effect of TDP-43 dysregulation on RNA processing by demonstrating a marked effect on the transcriptome [24]. Models that manipulate phase separation behaviour and aggregation of TDP-43 have also provided insight into the biophysical properties of the transition from biologically functional condensates to pathological states [23], [38]. Additionally, cytoplasmic mislocalisation of zebrafish TDP-43 (ubiquitously expressed) has demonstrated that cytoplasmic accumulation of the protein, rather than loss of nuclear TDP-43, has pronounced effects on metabolic function [32].

An important point of distinction of this study is the neuron-specific expression of hsa.TDP-43 in (dNLS, G294V and WT) and the grossly normal development of the motor system. This contrasts many other zebrafish ALS overexpression models (TDP-43 and other ALS-associated proteins), which display ubiquitous overexpression that generally is associated with significant axonal developmental abnormalities and impaired motor function at 2–3dpf [24], [54], [76]–[82]. The absence of early developmental defects in these models allows the assessment of non-developmental cellular changes and examination of mechanistic differences over time. To date, such longitudinal studies in zebrafish have been limited. One model, which overexpressed the ALS-causative hexanucleotide repeat mutation within *C9orf72,* reported an ageing phenotype characterised by reduced number of mature spinal cord motor neurons and muscle atrophy [58] and two overexpression models based on ALS-causative SOD1 mutations have characterised NMJ deficits, motor neuron loss and motor deficits in adult zebrafish [73], [83]. However, to the best of our knowledge, longitudinal analysis of the effects of TDP-43 dysregulation (other than survival) beyond 16 days [32] has not previously been reported in zebrafish. The gradual buildup of cytoplasmic TDP-43 and the formation of cytoplasmic puncta inclusions, in addition to the development of NMJ deficits and progressive loss of motor neurons with age, closely reflect critical features of human disease progression [84] and highlight the contribution of age-related changes to these pathologies. Such delayed-onset models are especially critical for investigating the role of loss- or gain-of-function mechanisms in ALS pathogenesis. Notably, hsa.TDP-43^dNLS^ mice show both nuclear loss of function and cytoplasmic gain of function and rapidly reached disease end stage within 6 weeks [33]. However, the hsa.TDP-43^dNLS^ zebrafish did reveal a much milder phenotype (∼50% survival at 13 months). We hypothesise that the nuclear retention of endogenous TDP-43 mitigates the pronounced loss-of-function effects seen in mice, thereby delaying the onset of severe phenotypes. This rescue effect is notable as it indicates that nuclear supplementation of certain variants of TDP-43 may be highly effective in delaying disease onset and progression. Furthermore, it makes the hsa.TDP-43^dNLS^ zebrafish platform particularly valuable for studying cytoplasmic gain-of-function mechanisms and to examine these mechanisms over extended timeframes.

While *in vitro* models provide an efficient means for axonal transport assessments, *in vitro* studies have generally shown poor correlation with *in vivo* analysis [85], highlighting the need for suitable *in vivo* models. This is particularly relevant as restoration of axonal transport deficits has been recognised as a potential therapeutic approach [52]. Here, we took advantage of a mCherry-tagged KIF5A reporter to assess active axonal transport in zebrafish axons. Indeed, the earliest phenotype identified in the hsa.TDP-43 zebrafish was a reduced axonal transport speed at 3 dpf in hsa.TDP-43^G294V^ zebrafish. This finding is in line with previous reports from TDP-43 models, including a *Drosophila* model [86] and TDP-43 mutant primary mouse cortical neurons [86], non-TDP-43 models of ALS including SOD1 mice [87], [88], and patient iPSC-derived motor neurons [86]. No defects in axonal transport were evident in hsa.TDP-43^dNLS^ zebrafish, suggesting that this pathology does not develop as a consequence of a TDP-43 cytoplasmic toxic gain of function in these fish. Further work is required to determine how the C-terminal point mutation G294V alters TDP-43 function specifically to affect axonal transport.

Our observation of an increase in the number of cytoplasmic TDP-43 puncta between 3 dpf and 6 dpf in all hsa.TDP-43 zebrafish may represent the very early stages of a liquid-to-solid transition of biomolecular condensates into insoluble aggregates. However, further investigation is required to assess the solubility and behavioural changes of these TDP-43 puncta over time. Notably, we did detect an increase in phosphorylated TDP-43 predominantly in the cytoplasm of aged (6 months) hsa.TDP-43 zebrafish motor neurons (**Fig 7**) which seems to reflect diffuse punctate cytoplasmic staining (DPCS) previously described as an early-stage pathology of ALS patients [89]. However, distinct pTDP-43 positive aggregates (skein-like, round or linear wisp shaped) which are a characteristic of advanced ALS [89] were not observed, conceivably suggesting that secondary insults are required for the progression of this pathology to occur - in line with the multi-hit hypothesis of neurodegenerative disease [90], [91]. Alternatively, in the examined samples, neurons expressing pTDP-43 aggregates may have already perished by 6 months of age.

Notably, hsa.TDP-43^dNLS^ zebrafish displayed a pronounced sensitivity to oxidative stress, highlighting the role of redox imbalance in exacerbating TDP-43 dysfunction associated with cytoplasmic accumulation of TDP-43. Consistent with this observation, we showed that Edaravone, an antioxidant, rescued motor deficits, highlighting the potential utility of the hsa.TDP-43 zebrafish in intervention studies. The differential response of the hsa.TDP-43 zebrafish to Riluzole, exacerbating motor dysfunction in hsa.TDP-43^WT^ zebrafish, highlights the need for further studies examining the mechanisms at play in ALS and the importance of model selection in therapeutic trials. Riluzole, which acts to reduce neuronal hyperexcitability, has been shown to reduce neuronal stress in SOD1 zebrafish models of ALS [92], a model that demonstrates early motor neuron and interneuron hyperexcitability. However, similar beneficial effects were not evident in a TDP-43 mouse model of ALS [93], consistent with our observations in the hsa.TDP-43 zebrafish. The lack of beneficial effect of Riluzole treatment in the hsa.TDP-43 zebrafish suggests that neuronal hyperexcitability may not be a prominent feature of this model of TDP-43 proteinopathies.

### Limitations

While zebrafish offer some unique advantages, they also do not fully reflect all TDP-43 pathologies, as evidenced by the absence of large pTDP-43 positive aggregates in the motor neurons of aged fish. The differences with mammalian systems must also be considered, and the interplay between TDP-43 gain-of-function toxicity and loss-of-function mechanisms, such as impaired cryptic exon repression, warrants further exploration.

## Conclusions

The progressive accumulation of cytoplasmic TDP-43 puncta and motor neuron loss in our model system mirrors key aspects of human ALS progression. Importantly, our results demonstrate that ageing exacerbates these phenotypes, with significant motor neuron loss observed in older zebrafish. Ageing is considered the primary risk factor influencing TDP-43 proteinopathies [94], and these hsa.TDP-43 models present an opportunity to study these effects *in vivo* over a prolonged timeframe. Patient studies are limited to end stage pathological observations and therefore require assumptions about the preceding events and while murine studies allow for snapshots of pathologies in time, repeated observations from the same animal are limited. This makes the hsa.TDP-43 zebrafish a valuable new tool to allow development of a temporal roadmap of TDP-43 pathology, from healthy to mislocalised and hyperphosphorylated states, and for testing drugs at specific pathological stages, with the ability to measure the effects of such interventions on individual animals or even neurons.

## Abbreviations

TDP-43: TAR DNA-binding protein 43
ALS: amyotrophic lateral sclerosis
FTLD: frontotemporal lobar dementia
LATE: limbic-predominant age-related TDP-43 encephalopathy
AD: alzheimer’s disease
dNLS: disrupted nuclear localisation sequence
STMN2: stathmin-2
UNC13A: unc-13 homologue A
LLPS: liquid-liquid phase separation
Hsa: TDP-43 human TDP-43
zTDP-43: zebrafish TDP-43
dpf: days post fertilisation
hpf: hours post fertilization
WT: wildtype
NonTg: Non-transgenic
NMJ: neuromuscular junction
mpf: months post fertilisation
SEM: standard error of the mean
EGFP: green fluorescent protein
N/C ratio: nuclear-cytoplasmic ratio
nTDP-43: native TDP-43 (not phosphorylated)
pTDP-43: phosphorylated-TDP-43

## Declarations

### Ethics approval

All experimental procedures were performed in compliance with the Macquarie University Animal Ethics Committee (ARA 2015/033, 2017/019) and the Macquarie University Biosafety Committee (NLRD 5974 – 52019597412350).

### Consent for publication

Not applicable

### Resource availability

The datasets used and/or analysed during the current study are available from the corresponding author on reasonable request.

### Author contributions

AH, MK, PC, GR, CM and SW performed the benchwork for the study. NS and ED helped to establish the hTDP-43 zebrafish lines. AH and MM conceived and designed the project. AH and MM wrote the manuscript with editing and intellectual input from ED, AL, IB, and RC. All authors reviewed and approved the final manuscript.

## Acknowledgements

We thank the Macquarie University Animal Services (MARS) staff for expert animal care, specifically Jason Martin-Powell and Cheryl Song for their daily animal support, Kelly Foskett and Tyler Chapman for their help in generating molecular constructs and Maxinne Watchon for her expertise designing the drug treatment trials.

