## Supplementary figures for "Human TDP-43 overexpression in zebrafish motor neurons triggers MND-like phenotypes through gain-of-function mechanism"

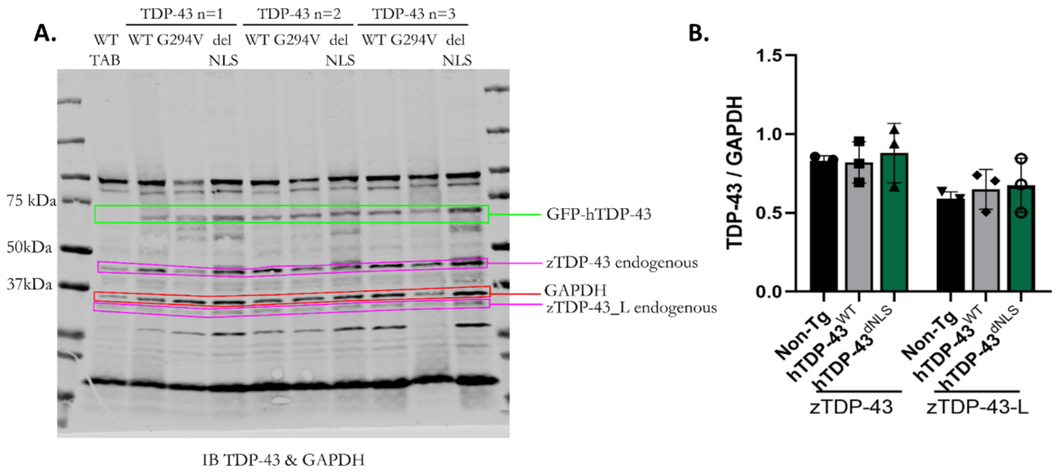


**Supplementary Figure 1 Western blot of hsa.TDP-43 zebrafish: A.** Uncropped western blot of lysates collected from *-3mnx1*:EGFP-hsa.TDP-43 zebrafish larvae at 3 dpf in triplicate. TDP-43 and GAPDH antibodies were used for analysis. **B.** Relative expression level of hsa.TDP-43 to endogenous zTDP-43. The hsa.TDP-43 transgene is expressed at a lower level than endogenous TDP-43 (hsa.TDP-43$^{WT}$ mean = 38%, hsa.TDP-43$^{G293V}$ at 59%, and hsa.TDP-43$^{dNLS}$ at 75%).

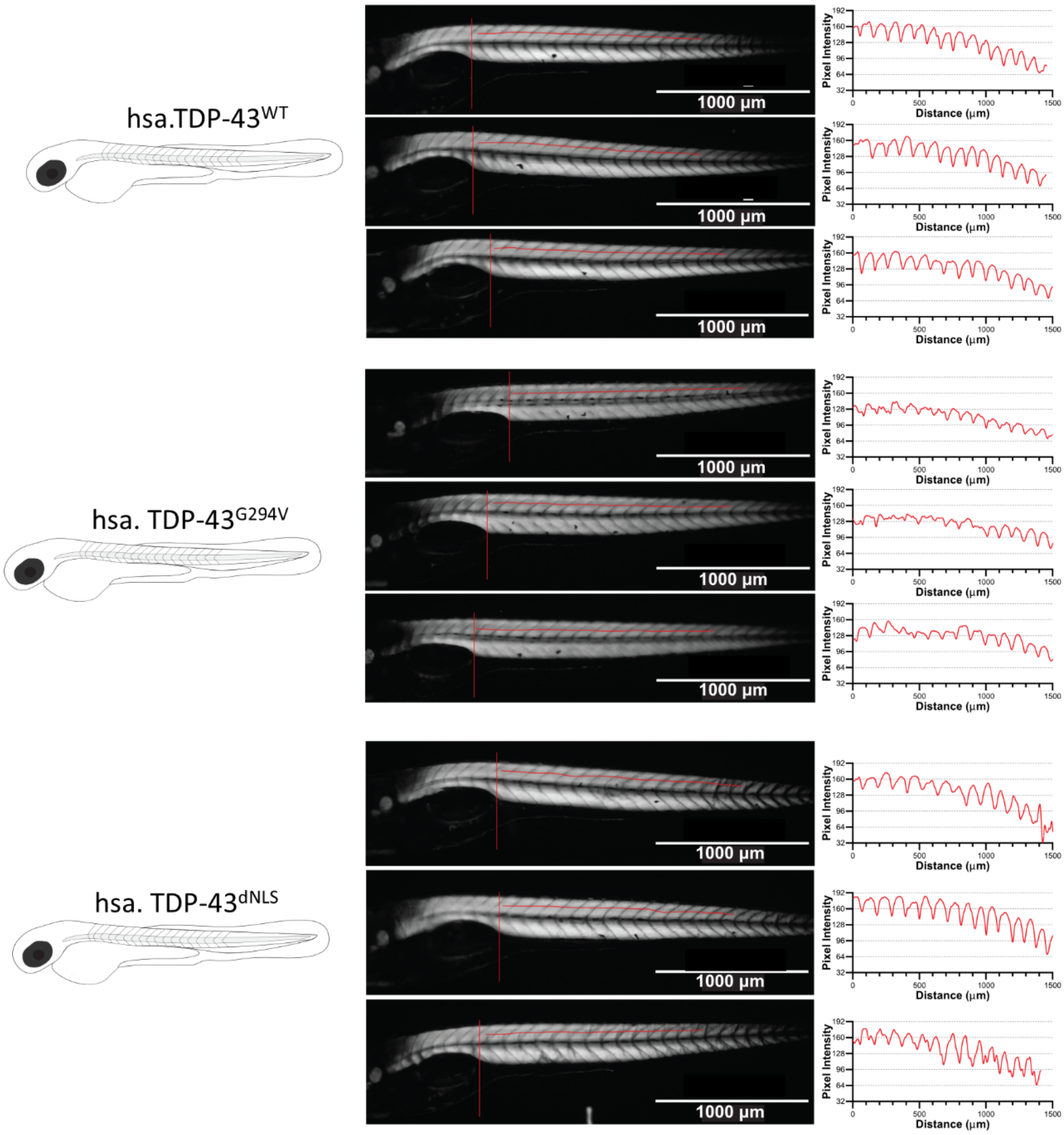


**Supplementary Fig 2**: **Birefringence analysis to examine muscle development in hsa.TDP-43 expressing zebrafish.** Analysis of birefringence at 6 dpf demonstrated no defects in myofibrillar alignment in the hsa.TDP-43 zebrafish, indicating no effect on muscle integrity (n = 3 fish per group).

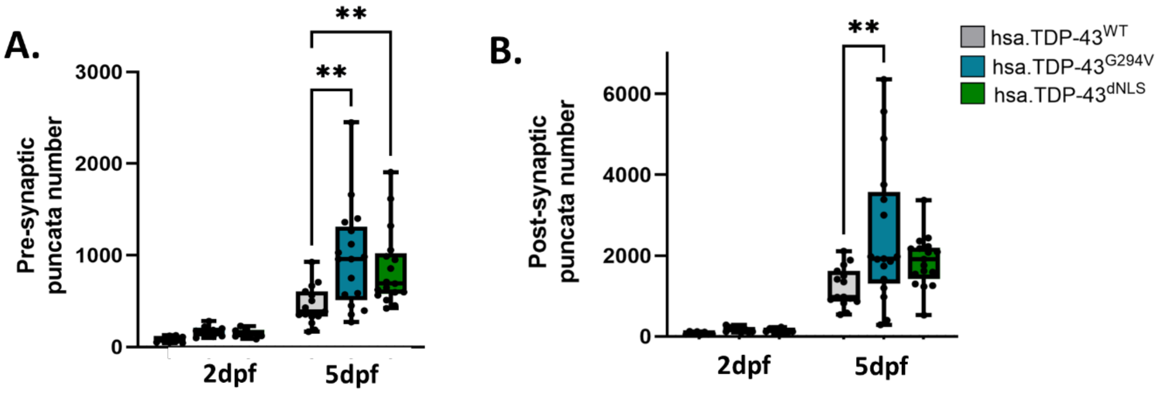


**Supplementary Figure 3: Quantification of the number of pre- and post- synaptic puncta at the neuromuscular junction. A.** No difference in the number of pre-synaptic puncta was evident between hsa.TDP-43 embryos at 2 dpf (hsa.TDP-43$^{WT}$ zebrafish mean = 80 +/- 12, hsa.TDP-43$^{G294V}$ zebrafish mean = 168 +/-20 and hsa.TDP-43$^{dNLS}$ zebrafish mean = 147 +/- 15). However, by 5 dpf, the number of pre-synaptic puncta in the hsa.TDP-43$^{G294V}$ zebrafish (mean = 975 +/- 135) and the hsa.TDP-43$^{dNLS}$ zebrafish (mean = 853 +/- 101) was significantly increased compared to hsa.TDP-43$^{WT}$ zebrafish (mean = 452 +/- 54, p=0.0019 and p = 0.0055 respectively) (One-way ANOVA). **B.** The number of post-synaptic puncta at 5 dpf was also significantly increased in the hsa.TDP-43$^{G294V}$ zebrafish (mean = 2507 +/-426) compared to hsa.TDP-43$^{WT}$ zebrafish (mean = 1217 +/- 125, p = 0.0066). The number of post-synaptic puncta was not significantly increased in hsa.TDP-43$^{dNLS}$ zebrafish relative to hTPD-43$^{WT}$ (mean = 1861 +/- 153) compared to hsa.TDP-43$^{WT}$ zebrafish (p = 0.25, One-way ANOVA).

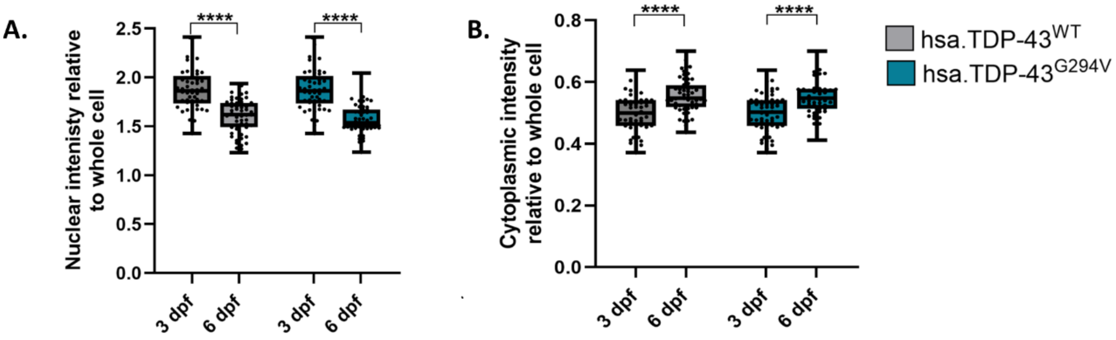


**Supplementary Figure 4. Reduced nuclear – cytoplasmic ratio from 3 dpf – 6 dpf is due to reduced nuclear hsa.TDP-43 expression and increased cytoplasmic expression. A.** The average nuclear fluorescence intensity of hsa.TDP-43 relative to whole cell average fluorescent intensity reduced between 3 dpf and 6 dpf in both hsa.TDP-43$^{WT}$ (1.9 +/- 0.03 to 1.6 +/- 0.2, p <0.0001) and hsa.TDP-43$^{G294V}$ (1.9 +/- 0.03 to 1.6 +/- 0.02, p <0.0001). **B.** The average cytoplasmic fluorescence intensity of hsa.TDP-43 relative to whole cell average fluorescent intensity increased between 3 dpf and 6 dpf in both hsa.TDP-43$^{WT}$ (0.5 +/- 0.008 to 0.6 +/- 0.008, p <0.0001) and hsa.TDP-43$^{G294V}$ (0.5 +/-0.008 to 0.55 +/- 0.009, p <0.0001).

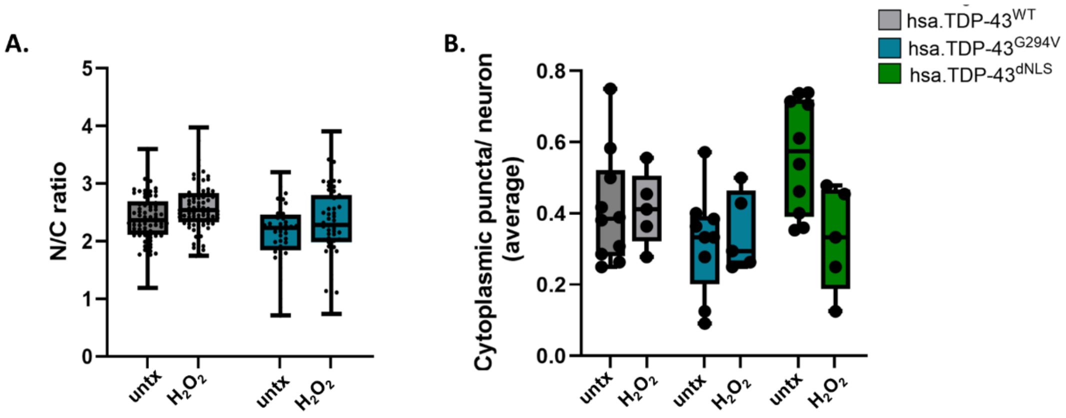


**Supplementary Figure 5.** Induction of oxidative stress by application of 0.5 mM $H_2O_2$ from 3 – 6 dpf did not affect localisation of hsa.TDP-43 or the incidence of cytoplasmic puncta formation. **A.** The nuclear-cytoplasmic ratio of hsa.TDP-43 at 6 dpf was not altered in primary motor neurons of treated compared to untreated hsa.TDP-43$^{WT}$ (mean = 2.4 +/- 0.05 v 2.6 +/- 0.05) or hsa.TDP-43$^{G294V}$ zebrafish (2.1+/- 0.08 v 2.3 +/- 0.09), indicating no cytoplasmic shift in the protein. **B.** The number of cytoplasmic puncta observed within primary motor neurons at 6 dpf was not increased in hsa.TDP-43$^{Wt}$ (mean = 0.41 +/- 0.05 v 0.41 +/- 0.046) hsa.TDP-43$^{G294V}$ (mean = 0.32 +/- 0.048 v mean = 0.35 +/- 0.5) or hsa.TDP-43$^{dNLS}$ zebrafish (mean = 0.56 +/- 0.05 v mean = 0.33 +/- 0.07).

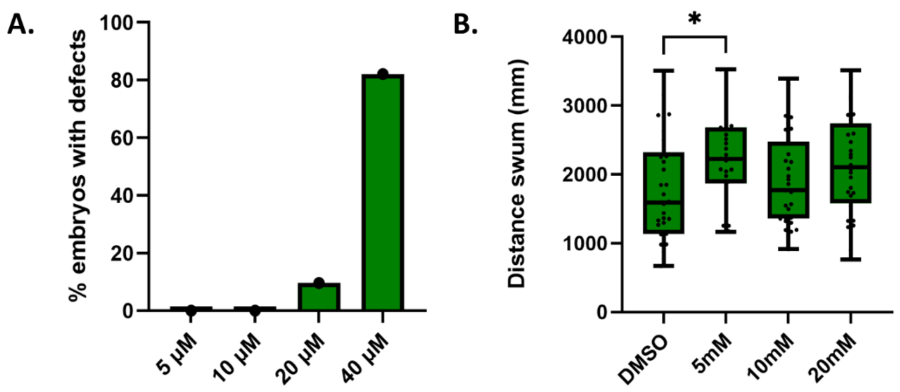


**Supplementary Fig 6: Optimisation of Edavarone treatment in hsa.TDP-43$^{dNLS}$ zebrafish. A.** Edavarone treatment at 5–100 μM was tested over 3 days to assess toxicity in the hsa.TDP-43$^{dNLS}$ zebrafish. Doses > 40 μM were lethal. Increased incidence of morphological abnormalities, primarily kinked tails and cardiac oedema were observed at 20 μM and 40 μM. **B.** Edavarone doses 5–20 μM were further assessed to determine their effect on motor function. 5 μM was found to be associated with a significant increase in distance swum by hsa.TDP-43$^{dNLS}$ zebrafish ($p = 0.04$) compared to DMSO-treated controls. Higher doses of 10 μM and 20 μM did not significantly improve motor function ($p = 0.7$ and $p = 0.1$ respectively). A 5 μM dose was therefore used in subsequent treatment trials.
